# Directional growth of developing myelin mediated by Wnt gradient and required for proper axon functions

**DOI:** 10.1101/2025.06.13.659097

**Authors:** Peng Liu, Qiu-Sui Deng, Qiang Chen, Xiang Chen, Jiu-Lin Du, Cheng He

**Author notes:** These authors contributed equally (Peng Liu, Qiu-Sui Deng). Correspondence (J.L.D.); (C.H.).

## Abstract

Axon-wrapping myelin sheaths formed by oligodendrocytes are essential for proper functions of the central nervous system. Although much is known about oligodendrocyte development, how the myelin dynamically forms remains unclear. Here we show the preferentially unidirectional extension of developing myelin mediated by Wnt gradient and required for proper axon functions. Using larval zebrafish as an *in vivo* model, we found that developing myelin in the spinal cord preferentially extends to the anterior end, and this process is dependent on an anterior-to-posterior Wnt4b gradient. Taking advantage of the large size of Mauthner-cell axons, we further showed that disruption of this directional extension impairs the even length distribution of myelin sheaths and faithful transduction of action potentials along the axon, and reduces the reliability of escape behavior. Thus, our study reveals a novel process for precise regulation of myelination, providing a new insight into myelin structuring and functioning.

## INTRODUCTION

In vertebrates, the unique compact myelin sheath is required for proper function of the central nervous system (CNS), facilitating rapid saltatory conduction of neural activity along wrapped axons, ensuring its long-term integrity, providing nutritional support, contributing to the functional regulation of nervous system plasticity, and playing an important role in learning and memory^1–5^. Myelin deficiencies result in not only impairment of nerve conduction but also axonal pathology and degeneration^6–8^. Thus, studies about the cellular and molecular mechanisms of myelin formation and maintenance are important for understanding basic elements of nervous system development and disease.

In the CNS, the transformation from membranous sheets elaborated by oligodendrocytes (OLs) into multi-layered compact myelin sheaths along axons is one of the most remarkable and complex developmental processes^9^. When OL-axon contacts are established, myelin segments grow via two different concerted motions - spiral wrapping around axons and lateral extension along axons^9^. Although the wrapping of the innermost layer of the growing myelin sheath has been well elucidated^10^, the dynamic lateral extension remains elusive^11^.

The larval zebrafish is an ideal system for studying myelination processes *in vivo*^12–14^. The basic features of myelin are conserved between mammals and zebrafish. Moreover, due to their embryonic transparency, zebrafish with fluorescent labelling in OLs enable the real-time visualization of morphogenetic development of myelin sheaths in vivo^15,16^. Here, we found the myelin sheaths exhibit a preference for anterior-orientated extension mediated by Wnt gradient in the spinal cord (SC) of zebrafish. Furthermore, disruption of this directional extension impairs the even length distribution of myelin sheaths and faithful transduction of action potentials (APs) on the Mauthner-cell axon (M-axon), and reduces the reliability of escape behavior. Thus, these findings reveal that the unidirectional extension of myelin sheath mediated by Wnt gradient acts as a novel cellular base for precise regulation of myelin formation and functioning during development.

## RESULTS

### Developing myelin preferentially extends to the anterior end along M-axons

The M-axon, with a large diameter, is a ventral projection axon from the hindbrain to the posterior end of the SC, providing an ideal model for visualizing the process of myelin development^15^. Individual OL and myelin segment can be mosaically labelled by mRFP under the control of the *Sox10* promoter^16^. By expressing *Sox10:mRFP* in the Tg(hspGFF62A,UAS:GFP) or Tol056 transgenic line, in both which GFP is expressed along M-axons (Figures 1A and S1A), we found that M-axons and myelin sheaths well co-localized in the ventral SC (Figures 1B and S1B). Moreover, by transient co-expression of *Sox10:eGFP* and *Sox10:mRFP* (Figure 1A), we found that the entrance point from oligodendroglial process to myelin segments was preferentially located on the posterior side of M-axon-wrapping myelin at late myelinating stage 6 days post-fertilization (dpf) (Figures 1C and 1E), implying the possibility of anterior extension of myelin sheaths along M-axons.

**Figure 1.**
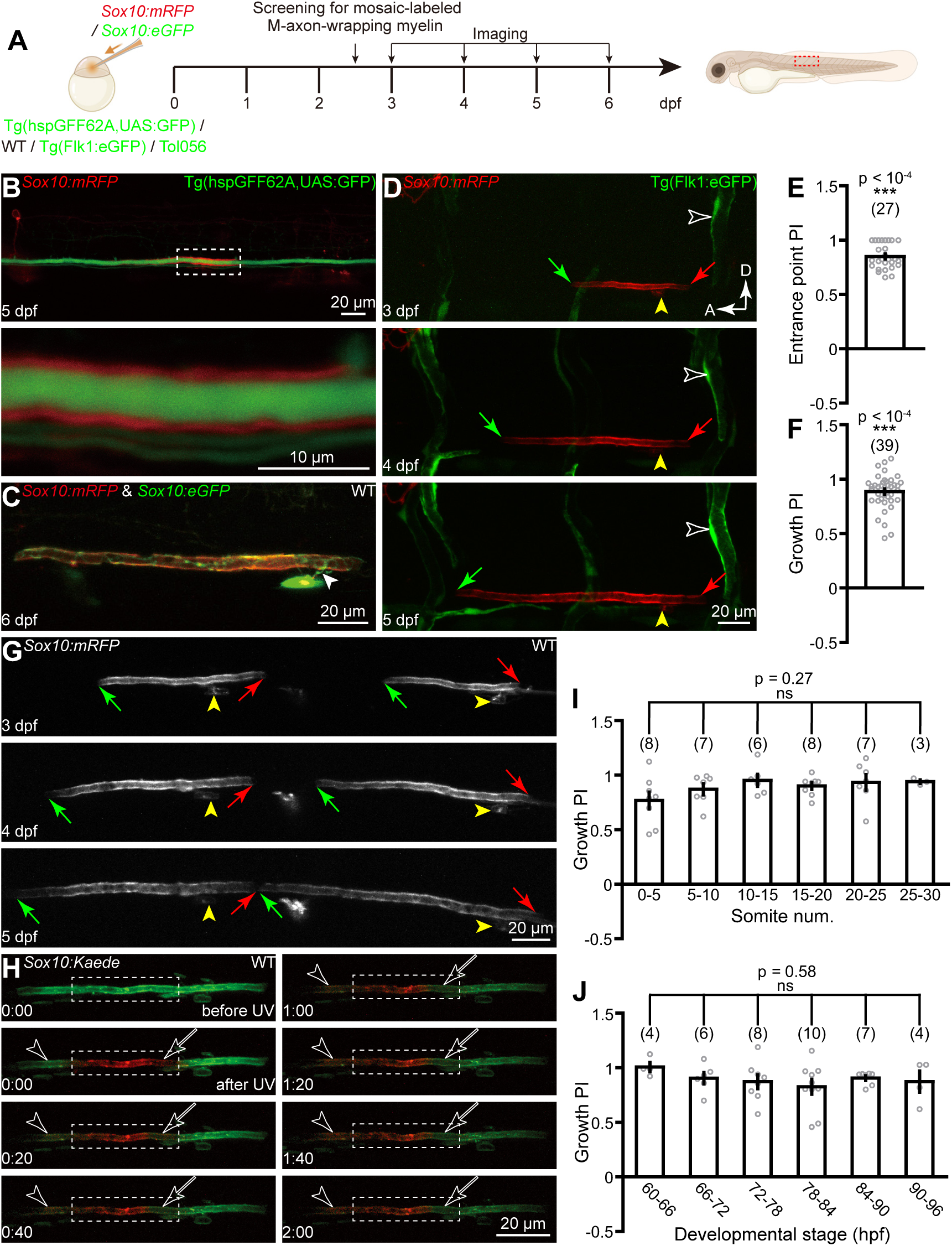
Anterior extension preference of M-axon-wrapping myelin sheaths during OL myelination *in vivo*. (A) Diagram of single M-axon-wrapping myelin sheath mosaically labeled by *Sox10:eGFP* and *Sox10:mRFP* transient expression in WT, Tg(hspGFF62A,UAS:GFP), Tg(Flk1:eGFP) and Tol056 lines. Timeline indicating the screening and imaging procedures. (B) Visualization of both myelin segments and M-axons. Top, projected image adopted from the red dotted frame in right cartoon fish in (A). Bottom, confocal image enlarged from the boxed area in top panel. Lateral views with dorsal up and anterior left. The same below unless noted otherwise. Dpf, days post-fertilization. (C) M-axon-wrapping myelin produced by an individual OL with cytoplasm in green and membrane in red acquired from larvae transiently expressing both *Sox10:eGFP* and *Sox10:mRFP*. Arrowhead, entrance point from oligodendroglial process to the M-axon-wrapping myelin. (D) Time-lapse images of M-axon-wrapping myelin segment in Tg(Flk1:eGFP) larvae. White hollow arrowhead, ISVs as reference markers; Yellow arrowhead, OL soma as fixed point of a reference marker during development; Green and red arrows, anterior and posterior ends of M-axon-wrapping myelin segment; D, dorsal; A, anterior. (E) PI of entrance point’s location along M-axon-wrapping myelin segments. PI is calculated as the subtraction of length from entrance point to anterior end and posterior end divided by the total length of myelin segment. Data were obtained from 27 OLs in 27 animals. (F) Growth PI of M-axon-wrapping myelin segments. Data were obtained from 39 OLs in 39 animals. (G) Time-lapse images of anterior extension of two neighboring M-axon-wrapping myelin segments. (H) Time-lapse images of Kaede-expressing M-axon-wrapping myelin with photoconversion in the middle region. Dotted box, UV irradiation region; Hollow arrowhead and arrow, anterior and posterior area of irradiation region. The timing before and after irradiation is depicted in the lower left. (I) Location-independency of Growth PI. Same data set with (F) was used. (J) Developing stage-independency of Growth PI. Same data set with (F) was used. The number on each column in (E), (F), (I) and (J) represents the number of M-axon-wrapping myelin segment used for statistics. Error bars, SEM. ns, not significant; **\*\*\***p < 0.001 (unpaired two-tailed Student’s t-test for (E) and (F) in comparison with 0; one-way ANOVA for (I) and (J)). See also Figures S1-S3.

Next, we performed *in vivo* long-term time-lapse confocal imaging of individual M-axon-wrapping OLs during 3 - 5 dpf with an interval of 12 hr, with different tissues and cellular organelles as reference markers to assess growth direction. We found that, after the first touch of M-axons by OL process, the myelin sheath exhibited a preference for anterior-orientated extension (Figures 1D and S1C-S1E), while intersegmental vessels (ISVs) in Tg(Flk1:eGFP) larvae (Figure 1D, hollow white arrowheads), OL somata (Figures S1C-S1E, yellow arrowheads) and nonspecific-labeled neurons (Figure S1C, hollow white arrowheads) all remained relatively immobile during development. The anterior and posterior ends of myelin segments at each time-point were confirmed by slice-to-slice characterization of Z-stack images (Figure S2). We used growth preference index (Growth PI), which was calculated as the subtraction of extension length in anterior and posterior ends divided by total extension length, to characterize the directional extension preference of myelin sheaths. Positive values indicate anterior extension preference. The average Growth PI was 0.88 ± 0.03 (Figure 1F), and it did not change with location (Figure 1I) or developmental stage (Figure 1J). In the following experiments, OL somata were mainly used as reference to assess myelin growth direction.

We then monitored the developmental procession of two neighboring myelin segments along the M-axon and found both the two myelin sheaths showed anterior-orientated extension (Figures 1G and S3). Furthermore, to examine growth direction at the level of myelin membrane movement, we expressed the photoconvertible fluorescent protein Kaede to precisely label membrane component in the middle area of a single myelin segment under ultraviolet (UV) irradiation (Figure 1H, dotted boxes). During two hours after irradiation, the anterior region of the photoconvertible area turned into red from green (Figure 1H, white arrowheads), while the posterior region turned into green from red (Figure 1H, white arrows), indicating an anterior movement of myelin membrane.

### Wnt4b signaling is required for the anterior extension of M-axon-wrapping myelin

In the SC, morphogens, including Fibroblast Growth Factor (FGF), Sonic Hedgehog (Shh), Bone Morphogenetic Protein (BMP) and Wnt signaling, govern the pattern of tissue development and the position of various specialized cell types during embryogenesis^17–22^. To exam whether such morphogens contribute to the directional growth of myelin segments, we employed specific antagonist to block their signaling pathways (Figure 2A). Our results demonstrated that Wnt signaling inhibition significantly impaired the growth preference of M-axon-wrapping myelin sheathes, whereas other morphogen antagonist showed no detectable effects (Figures 2B-2G). These results suggest that Wnt-mediated signaling plays an important role in orchestrating the spatial regulation of myelination.

**Figure 2.**
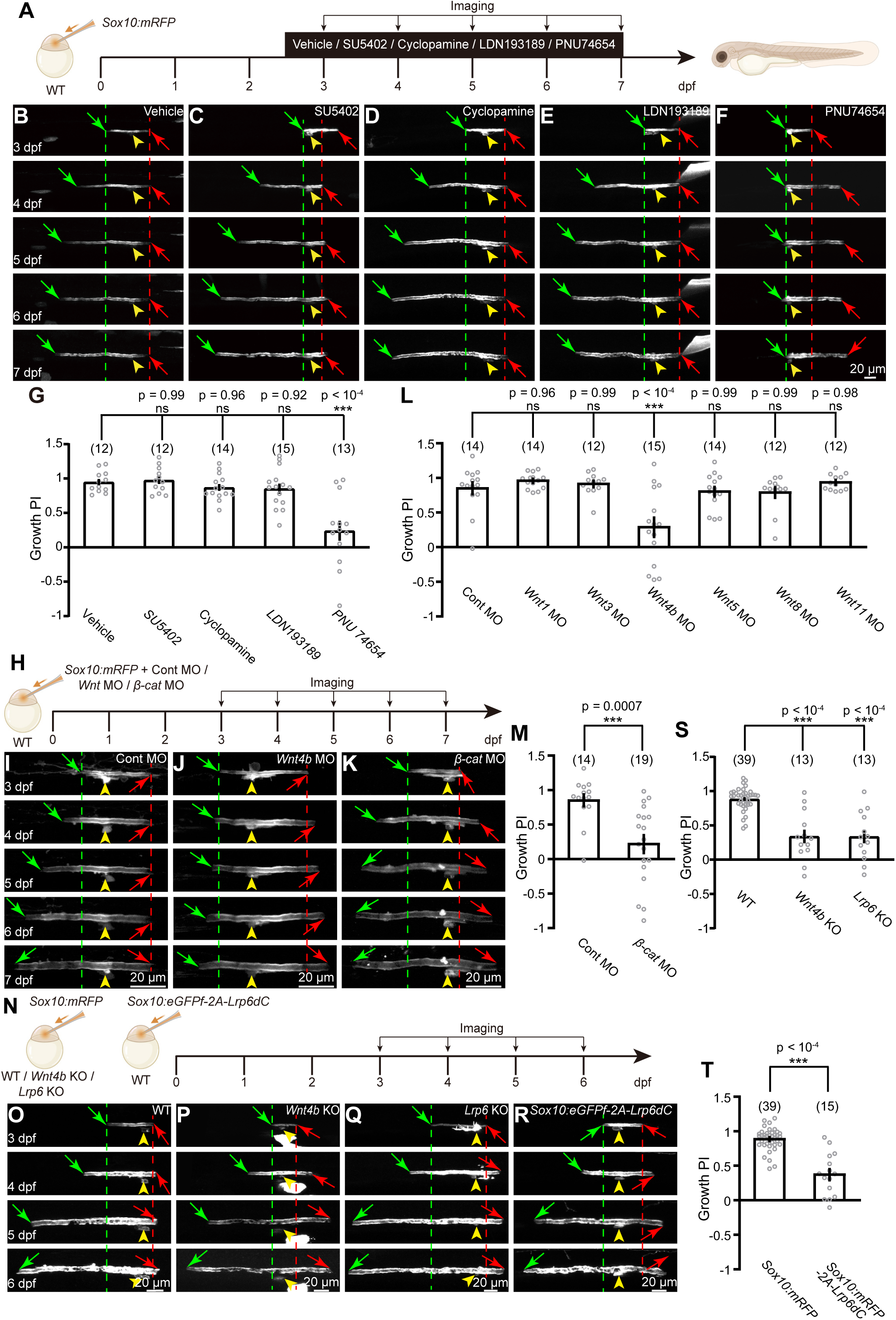
Oligodendroglial Wnt signaling is required for the unidirectional preference of developmental myelin extension. (A) Schematic diagram displaying the time course for morphogen antagonist treatment. Timeline indicating the imaging procedures. (B-F) Time-lapse images of M-axon-wrapping myelin with different morphogen antagonist treatment. (B) control. (C) SU5402, an FGF inhibitor. (D) Cyclopamine, Sonic Hedgehog signaling inhibitor. (E) LDNf193189, BMP pathway inhibitor. (F) PNU74654, Wnt/β-catenin pathway inhibitor. Green and red lines indicate the anterior and posterior ends of M-axon-wrapping myelin segment at 3 dpf. (G) Growth PI of M-axon-wrapping myelin segments with vehicle and morphogen antagonist treatment. (H) Schematic diagram displaying the imaging timeline of M-axon-wrapping myelin in control, *Wnt* and *β-catenin* morphants. (I-K) Time-lapse images of M-axon-wrapping myelin in control (I), *Wnt4b* (J), and *β-catenin* (K) morphants. (L) Growth PI of M-axon-wrapping myelin segments in control and *Wnt* morphants. (M) Growth PI of M-axon-wrapping myelin segments in control and *β-catenin* morphants. Same data set with (L) was used in Control MO. (N) Diagram of single M-axon-wrapping myelin sheath mosaically labeled by *Sox10:mRFP* transient expression in WT, *Wnt4b* KO and *Lrp6* KO lines, and *Sox10:eGFPf-2A-Lrp6dC* in WT. Timeline indicating the imaging procedures. (O-Q) Time-lapse images of M-axon-wrapping myelin in WT (O), *Wnt4b* KO(P), and *Lrp6* KO (Q) lines. (Q) Time-lapse images of M-axon-wrapping myelin labeled by *Sox10:eGFPf-2A-Lrp6dC* transient expression. (R) Growth PI of M-axon-wrapping myelin segments in WT, *Wnt4b* KO, and *Lrp6* KO lines. Same data set with Figure 1F was used in WT larvae. (S) Growth PI of M-axon-wrapping myelin segments in *Sox10:mRFP* and *Sox10:mRFP-2A-Lrp6dC* larvae. Same data set with Figure 1F was used in *Sox10:mRFP* larvae. The number on each column in (G, L, M, S and T) represents the number of M-axon wrapping myelin segments examined. Error bars, SEM. ns, not significant; **\***p < 0.05, **\*\***p < 0.01, **\*\*\***p < 0.001 (unpaired two-tailed Student’s t-test for (M) and (T); one-way ANOVA for (G), (L) and (S)). See also Figures S4 and S5.

To further dissect the function of Wnt signaling, we used previously characterized morpholino oligomers (MOs) to knock down the expression of Wnt ligands in zebrafish larvae, including Wnt1, Wnt3, Wnt4b, Wnt5, Wnt8 and Wnt11. The knockdown efficiency was confirmed by reduced fluorescence intensity in the Wnt reporter line Tg(TCF:GFP) (Figures S4A, S4B). The results showed that only in *Wnt4b* morphants, the directional preference was markedly impaired (Figures 2L, 2H, 2I, and 2J). Moreover, by knocking down *β-catenin* expression, a key molecule at downstream of Wnt receptors, we found that the extension preference was also significantly disrupted (Figures 2K and 2M). Intriguingly, we found that Wnt4b, but not β-catenin, significantly impaired the dorsal migration of oligodendrocyte lineage cells in the SC of zebrafish at 3 dpf (Figures S4C and S4D).

Given the embryonic lethality observed in β-catenin knockout fish, we further generated frame-shift mutant zebrafish by CRISPR-Cas9 technology with 22 and 13 bases ablation of *Wnt4b* and *Lrp6* gene, the co-receptor of Wnt signaling, respectively (Figures S5A-S5F). We found that both Wnt4b and Lrp6 KO fish showed significantly reduced growth preference indices (GPI) of myelin sheaths compared to controls (Figures 2O–2Q, 2S). At the same time, the myelinating abilities, such as OPC migration and myelin formation, were severely impaired (Figures S5G-S5J).

Furthermore, we specifically expressed Lrp6dC, a dominant interfering mutant of the Wnt co-receptor LRP6, in OLs to block oligodendroglial Wnt signaling^23^. As a result, the extension preference was also markedly impaired (Figures 2R and 2T). The Tg(sox10:mRFP-Lrp6dC) line also exhibited compromised OL dorsal migration and myelination (Figures S4F-S4I).

Collectively, these results suggest the critical role of Wnt4b signaling in orchestrating the directional growth of myelinating sheaths.

### Wnt4b gradient is necessary for the anterior extension preference of M-axon-wrapping myelin

Previous studies have reported an anterior-to-posterior Wnt4 decreasing gradient in zebrafish and mammalian SC^24,25^. To further exam the role of Wnt4b gradients in regulating the directional growth of myelin sheaths, we first analyzed the expression pattern of *Wnt4b* in the SC by *in situ* hybridization. We found that *Wnt4b* enriched in the ventral SC, and exhibited an anterior-to-posterior decreasing gradient along the entire length of the floorplate (Figure 3A). It also showed a similar gradient in the cross sections at 3 dpf (Figures 3B and 3C).

**Figure 3.**
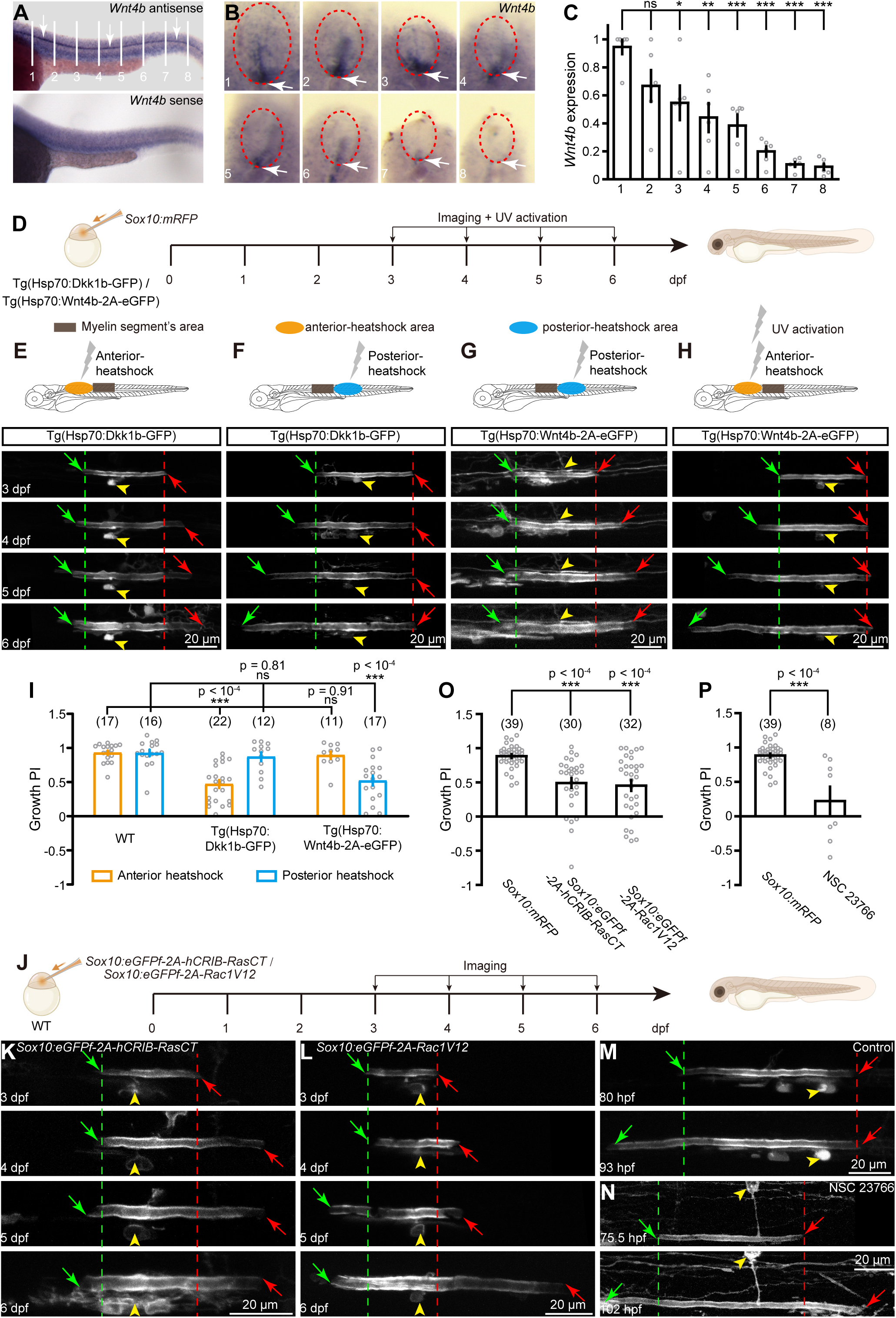
Wnt4b gradient is necessary for the anterior extension preference of M-axon-wrapping myelin. (A) Lateral view of *Wnt4b* expression pattern at 3 dpf by whole-mount *in situ* hybridization. White lines and numbers indicate the position of cross sections in (B). Arrowheads, the *Wnt4b* expression in the ventral SC. (B) *Wnt4b* expression pattern in the cross section of the SC. Oval frames, SC; Arrowheads, the *Wnt4b* expression in the ventral SC. (C) Anterior-to-posterior *Wnt4b* gradient in the SC. Relative *Wnt4b* expression intensity along anterior-to-posterior axis, normalized to the baseline level (section 1). Numbers indicate the position in (B). Data in the were obtained from 6 embryos. ns p = 0.22, **\***p = 0.018, **\*\***p = 0.0012, **\*\*\***p < 0.0001. (D) Diagram of single M-axon-wrapping myelin sheath mosaically labeled by *Sox10:mRFP* transient expression in Tg(Hsp70:Dkk1b-GFP) and Tg(Hsp70:Wnt4b-2A-eGFP) transgenic lines. Timeline indicating the imaging and UV activation procedures. (E-H) Time-lapse images of M-axon-wrapping myelin segments in Tg(Hsp70:Dkk1b-GFP) (E and F) and Tg(Hsp70:Wnt4b-2A-eGFP) (G and H) larvae with local infrared heat application. Square frame in the cartoon (top) indicates the region of myelin segments observed. Oval color frames indicate the local areas where infrared heat was applied. (I) Growth PI of M-axon-wrapping myelin in WT, Tg(Hsp70:Dkk1b-GFP) and Tg(Hsp70:Wnt4b-2A-eGFP) larvae with local infrared heat application. (J) Diagram of single M-axon-wrapping myelin sheath mosaically labeled by *Sox10:eGFPf-2A-hCRIB-RasCT* and *Sox10:eGFPf-2A-Rac1V12* transient expression. (K and L) Time-lapse images of M-axon-wrapping myelin labeled by *Sox10:eGFPf-2A-hCRIB-RasCT* (K) and *Sox10:eGFPf-2A-Rac1V12* (L) transient expression. (M and N) Time-lapse images of of M-axon-wrapping myelin with control (B) and Rac1 antagonist NSC23766 (150 μM) treatment. (O) Growth PI in *Sox10:mRFP*, *Sox10:eGFPf-2A-hCRIB-RasCT* and *Sox10:eGFPf-2A-Rac1V12* larvae. Same data set with Figure 1F was used in *Sox10:mRFP* larvae. (P) Myelin Growth PI in control and Rac1 antagonist treated larvae. Same data set with Figure 1F was used in *Sox10:mRFP* larvae. The number on each column in (I, O and P) represents the number of M-axon wrapping myelin segments examined. Error bars, SEM. ns, not significant; **\*\*\***p < 0.001 (two-way ANOVA for (I); one-way ANOVA for (C) and (O); unpaired two-tailed Student’s t-test for (P)).

To examine the possible role of the Wnt4b gradient in myelin anterior extension, we then suppressed the endogenous Wnt gradient by up-regulating the expression of the Wnt inhibitor Dkk1b^26^ in the rostral side of one myelin segment via local infrared heat application in Tg(Hsp70:Dkk1b-GFP) larvae^27,28^ (Figures 3D and 3E). We found that the Growth PI was significantly reduced (Figure 3I). Meanwhile, the anterior preference was not markedly affected by Dkk1b over-expression in the caudal side (Figures 3F and 3I), which was supposed to enhance the Wnt gradient. Furthermore, we weakened the endogenous Wnt gradient by local heat-induced Wnt4b overexpression in the caudal side of myelin segments in Tg(Hsp70:Wnt4b-2A-eGFP) larvae (Figures 3D and 3G), and found that myelin segments showed a decreased Growth PI (Figure 3I). But rostral Wnt4b overexpression had no significant effect (Figures 3H and 3I). Taken together, these results indicate that the anterior-to-posterior Wnt4b gradient is necessary and sufficient for the directional extension of myelin.

The small Rho GTPase Rac1, a Wnt downstream signal, is necessary for formation of membrane protrusion and migration of many types of cells^29^. To examine whether Rac1 activity is required for the directional extension of myelin, we used *Sox10* promoter-driven expression of the Rac1 inhibitor hCRIB-RasCT^30^ to impair endogenous Rac1 function in OLs (*Sox10:eGFPf-2A-hCRIB-RasCT*) (Figure 3J). We found that myelin segments displayed a significant reduction in the Growth PI (Figures 3K and 3O). Furthermore, myelin segments expressing constitutively active Rac1 (*Sox10:eGFPf-2A-Rac1V12*)^30^ also exhibited a lower PI (Figures 3L and 3O). Consistently, a similar phenomenon was observed by the treatment of the Rac1 antagonist NSC23766^31^ (Figures 3M, 3N and 3P).

### Preferentially anterior extension of non-M-axon-wrapping myelin sheaths is mediated by Wnt gradient in the ventral SC

To examine whether myelin directional extension is a general phenomenon or only specific to M-axons, we then performed time-lapse imaging of non-M-axon-wrapping myelin segments (Figure 4A). We found that, although extending to both anterior and posterior directions simultaneously, non-M-axon-wrapping myelin segments still showed an anterior preference in the ventral SC (Figures 4B and 4F). However, in the dorsal part, non-M-axon-wrapping myelin segment rarely exhibited a directional preference (Figures 4C and 4F). Interestingly, the *Wnt4b* expression showed a ventral-to-dorsal decreasing gradient in the SC (Figures 4F and 3B). Quantitative analysis revealed that the Growth PI was positively correlated with the level of *Wnt4b* expression (Figure 4F), with a strong anterior extension preference in the ventral SC where high levels of *Wnt4b* expressed. Moreover, the non-M-axon-wrapping myelin displayed a significant reduction of the Growth PI in ventral but not dorsal SC in *Sox10:mRFP-2A-Lrp6dC* larvae (Figures 4D, 4E and 4G). Furthermore, when an opposite Wnt gradient was established with local heat-induced overexpression of Wnt4b on the caudal side in Tg(Hsp70:Wnt4b-2A-eGFP) larvae (Figure 4H), myelin segments exhibited a reverse growth directional preference in particular in the ventral SC (Figures 4I-4K). These results suggest that the preferentially anterior extension of non-M-axon-wrapping myelin sheaths is mediated by Wnt4b gradient as well.

**Figure 4.**
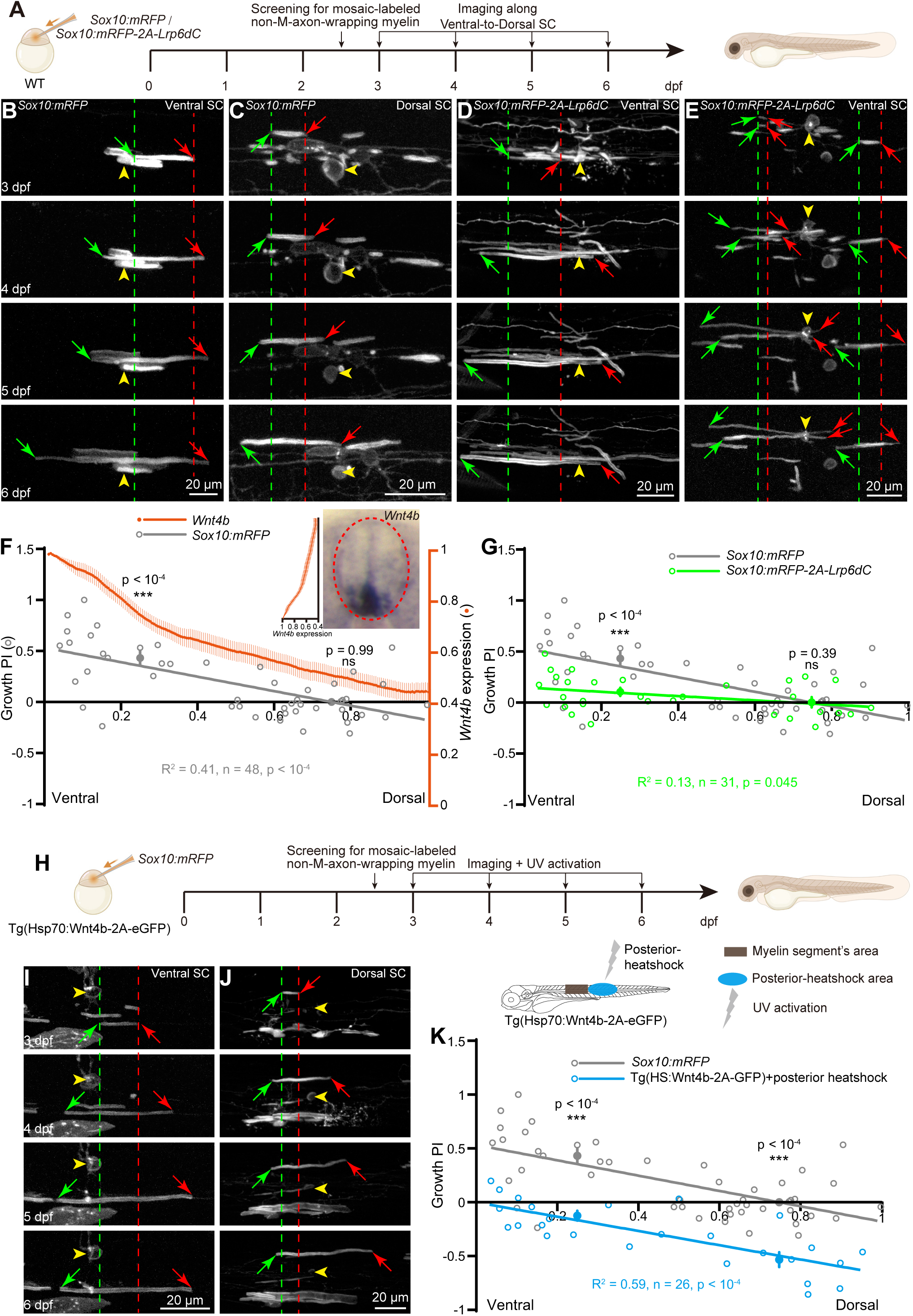
Anterior extension preference of non-M-axon-wrapping myelin mediated by Wnt gradient. (A) Diagram of single non-M-axon-wrapping myelin sheath mosaically labeled by *Sox10:mRFP* and *Sox10:mRFP-2A-Lrp6dC* transient expression. Timeline indicating the screening and imaging procedures. (B and C) Time-lapse images of non-M-axon-wrapping myelin in the ventral (B) and dorsal (C) SC of *Sox10:mRFP* larvae. Most of the myelin sheaths were taken from the middle region along anterior-to-posterior axis. (D and E) Time-lapse images of non-M-axon-wrapping myelin in ventral (D) and dorsal (E) SC of *Sox10:mRFP-2A-Lrp6dC* larvae. (F) Positive correlation between Growth PI of non-M-axon-wrapping myelin (grey) and *Wnt4b* expression intensity (orange) along ventral-to-dorsal axis in the SC of *Sox10:mRFP* larvae. The numbers of X axis indicate the relative positions along ventral (left) to dorsal (right) axis of the spinal cord. Inset panel, *Wnt4b* expression pattern in the cross section of SC. Oval frame, SC; Inset diagram, relative *Wnt4b* expression intensity (orange) along ventral-to-dorsal axis. Grey line, linear regression of myelin Growth PI along ventral-to-dorsal axis. Solid grey dots indicate the average Growth PI in the ventral and dorsal SC. (G) Linear regression of the Growth PI of non-M-axon-wrapping myelin along ventral-to-dorsal axis in the SC of *Sox10:mRFP* (grey) and *Sox10:mRFP-2A-Lrp6dC* (green) larvae. Solid green dots indicate the average Growth PI in the ventral and dorsal SC of *Sox10:mRFP-2A-Lrp6dC* larvae. Same data set with (F) was used for *Sox10:mRFP* larvae. (H) Diagram of single non-M-axon-wrapping myelin sheath mosaically labeled by *Sox10:mRFP* transient expression in Tg(Hsp70:Wnt4b-2A-eGFP) transgenic lines. Timeline indicating the imaging and UV activation procedures. (I and J) Time-lapse images of non-M-axon-wrapping myelin in ventral (I) and dorsal (J) SC of Tg(Hsp70:Wnt4b-2A-eGFP) larvae with local infrared heat application at the caudal side of the myelin segments. (K) Linear regression of the Growth PI of non-M-axon-wrapping myelin along ventral-to-dorsal axis in the SC of *Sox10:mRFP* (grey) and Tg(Hsp70:Wnt4b-2A-eGFP) with caudal side Wnt activation (blue) larvae. Solid blue dots indicate the average Growth PI in the ventral and dorsal SC of Tg(Hsp70:Wnt4b-2A-eGFP) larvae with Wnt activation in the caudal side. Same data set with (F) was used in *Sox10:mRFP* larvae. “n” represents the number of non-M-axon wrapping myelin used for statistics in (F), (G) and (K). Error bars, SEM. ns, not significant; **\*\*\***p < 0.001 (two-way ANOVA for (F) in comparison with 0; two-way ANOVA for (G) and (K) in comparison with the Growth PI in *Sox10:mRFP* larvae).

### Disruption of anterior extension preference impairs the uniformity of developing myelin sheaths

To explore the function of myelin anterior extension preference, we first characterized the process of oligodendroglial development in the SC. In vivo imaging of Tg(Olig2:eGFP) larvae, in which oligodendroglial linage cells express eGFP, demonstrated that oligodendrocyte precursor cells (OPCs) migration exhibited a sequential spatiotemporal pattern along the anterior-to-posterior axis (Figures S6A, S6C and S6H). Characterization of larva transiently expressing both *Sox10:mRFP* and *Sox10:eGFP* also revealed a similar pattern in oligodendroglial differentiation and maturation (Figures S6B, S6D and S6I). Consistent with this myelinating gradient, we found that myelin length decreased dramatically from the rostral to caudal side of the SC at 3 dpf, a time point around the myelinating initiation stage (Figures S6E and S6J). However, at 7 dpf, the late stage of myelination, there was no significant difference in myelin length along the anterior-to-posterior axis (Figures 5A, 5B and 5G), suggesting a process of uniformization of myelin segment length during development.

**Figure 5.**
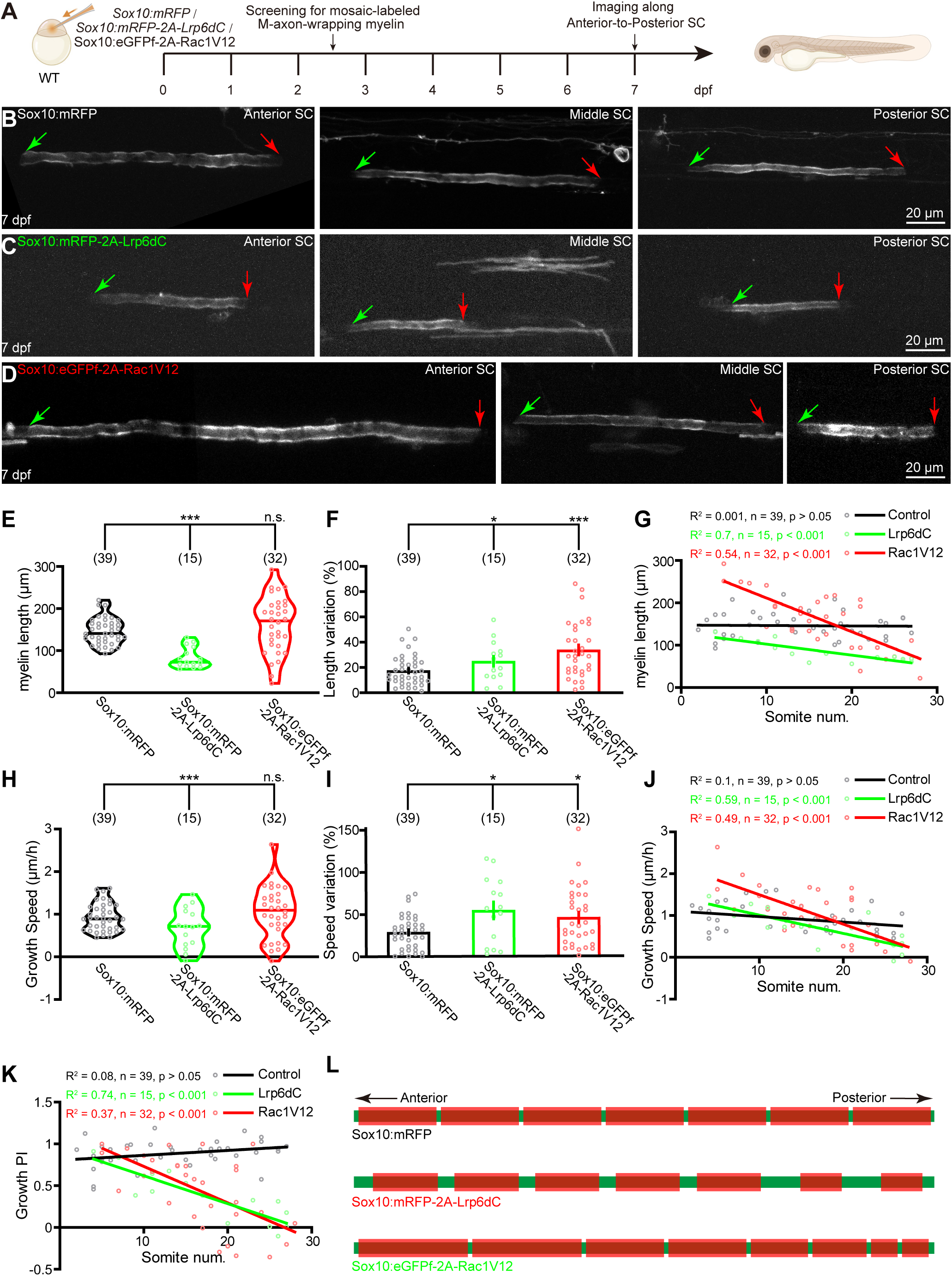
Even length distribution of myelin sheaths contributed by anterior extension preference. (A) Diagram of single M-axon-wrapping myelin sheath mosaically labeled by *Sox10:mRFP, Sox10:mRFP-2A-Lrp6dC* and *Sox10:eGFPf-2A-Rac1V12* transient expression. Timeline indicating the imaging procedures. (B-D) Confocal images of M-axon-wrapping myelin along anterior-to-posterior axis of the SC in *Sox10:mRFP* (B), and *Sox10:mRFP-2A-Lrp6dC* (C) and *Sox10:eGFPf-2A-Rac1V12* (D) larvae at 7 dpf. (E) Length of M-axon-wrapping myelin in *Sox10:mRFP*, *Sox10:mRFP-2A-Lrp6dC* and *Sox10:eGFPf-2A-Rac1V12* larvae at 7 dpf. Same data set with Figures 1F and 3O was used. (F) Length variation of M-axon-wrapping myelin in *Sox10:mRFP*, *Sox10:mRFP-2A-Lrp6dC* and *Sox10:eGFPf-2A-Rac1V12* larvae at 7 dpf. Same data set with (E) was used. (G) Linear regression of the length of M-axon-wrapping myelin distributed along anterior-to-posterior axis of the SC in *Sox10:mRFP* (black), *Sox10:mRFP-2A-Lrp6dC* (green) and *Sox10:eGFPf-2A-Rac1V12* (red) larvae at 7 dpf. Same data set with (E) was used. (H) Growth speed of M-axon-wrapping myelin in *Sox10:mRFP*, *Sox10:mRFP-2A-Lrp6dC* and *Sox10:eGFPf-2A-Rac1V12* larvae at 7 dpf. Same data set with (E) was used. (I) Growth speed variation of M-axon-wrapping myelin in *Sox10:mRFP*, *Sox10:mRFP-2A-Lrp6dC* and *Sox10:eGFPf-2A-Rac1V12* larvae at 7 dpf. Same data set with (E) was used. (J) Linear regression of the growth speed of M-axon-wrapping myelin distributed along anterior-to-posterior axis of the SC in *Sox10:mRFP* (black), *Sox10:mRFP-2A-Lrp6dC* (green) and *Sox10:eGFPf-2A-Rac1V12* (red) larvae at 7 dpf. Same data set with (E) was used. (K) Linear regression of the growth PI of M-axon-wrapping myelin distributed along anterior-to-posterior axis of the SC in *Sox10:mRFP* (black), *Sox10:mRFP-2A-Lrp6dC* (green) and *Sox10:eGFPf-2A-Rac1V12* (red) larvae at 7 dpf. Same data set with (E) was used. (L) Schematic of the length of M-axon-wrapping myelin distributed along anterior-to-posterior axis of the SC in *Sox10:mRFP* (left) and *Sox10:eGFPf-2A-Rac1V12* (right) larvae. A, anterior; P, posterior; Green frames, M-axons; Red frames, M-axon-wrapping myelin segments. The number on each column in (E-K) represents the number of M-axon wrapping myelin segments used for statistics. Error bars, SEM. ns, not significant; **\***p < 0.05, **\*\***p < 0.01, **\*\*\***p < 0.001 (two-way ANOVA for (E-K)). See also Figure S6.

To test whether the directional growth of myelin segments contributes to their length uniformity, we examined myelin length distribution in Sox10:mRFP-2A-Lrp6dC and Sox10:eGFPf-2A-Rac1V12 animals, in which oligodendrocyte-specific blockage of Wnt and downstream signaling impairs the anterior extension preference. We found that, at 3 dpf, both lines showed a rostral-to-caudal reduction of myelin segment length along the SC, similar with controls (Figures S6F, S6G and S6J). However, by 7 dpf, further analyses demonstrated a decreasing gradient in myelin length along anterior-to-posterior axis, with longer in rostral and shorter in caudal side of the SC in Sox10:eGFPf-2A-Rac1V12 and Sox10:mRFP-2A-Lrp6dC animals (Figures 5C, 5D and 5G), in comparison with the even length distribution in control, indicating that disruption of extension preference impairs the process of uniformization of myelin length. At the same time, the average length of myelin segments was comparable in Sox10:eGFPf-2A-Rac1V12 but decreased significantly in Sox10:mRFP-2A-Lrp6dC (Figure 5E), and the length variation increased remarkably in both animals (Figure 5F), consistent with the impairment of length uniformity. The growth speed of myelin segments demonstrated similar changes (Figures 5H-5J). Additionally, the growth PI also displayed a rostral-to-caudal decreasing gradient in both transgenic lines (Figure 5K). Thus, these results demonstrated that the directional extension preference of myelin segments contributes to the uniformity of their length distribution (Figure 5L).

Moreover, our results also showed that, in addition to length uniformity, the growth ability of myelin segments was impaired significantly in Sox10:mRFP-2A-Lrp6dC but not in Sox10:eGFPf-2A-Rac1V12 lines (Figures 5E and 5H). Further morphological examination demonstrated that dorsal migration and myelination of oligodendrocyte linage cells were comparable in Sox10:eGFPf-2A-Rac1V12 (Figures S6K-S6N), but significantly impaired in Sox10:mRFP-2A-Lrp6dC lines (Figures S4F-S4I). Consistent with these findings, TEM analysis showed a significant hypomyelination of M-axons in Tg(Sox10:mRFP-2A-Lrp6dC) but not Tg(Sox10:eGFPf-2A-Rac1V12) transgenic lines (Figures S4J-S4L). Thus, in the following function test of myelin directional extension, to avoid the effects of myelination deficits in the *Wnt4b* KO, *Lrp6* KO, and Tg(Sox10:mRFP-2A-Lrp6dC) lines, we employed Tg(Sox10:eGFPf-2A-Rac1V12) to investigate how myelin preferentially extension influences axonal function and behavior (Figure 5L).

### Deficiency in myelin preferentially extension impairs proper axon functions

To study the function of the directional extension of myelin at physiological levels, we assessed the conduction of APs along M-axons by *in vivo* whole-cell recording. Electrical stimulation of M-axon terminals evoked reliable APs that could be recorded from M-cell somata at 7 dpf (Figure 6A). We detected no overt difference in axon excitability between Tg(Sox10:eGFPf-2A-Rac1V12) and Tg(Sox10:mRFP) larvae (Figure S7A, left), confirming the overall integrity of myelinated tracts. Furthermore, evoked APs could not be suppressed by co-application of the AMPA receptor antagonist CNQX and the NMDA receptor antagonist D-AP5 (Figure S7A, middle), whereas vanished when the stimulating electrode was far away from the M-axon terminal (Figure S7A, right), implying the recorded APs were specifically evoked by the electrical stimulation on M-axons and propagated backward along the M-axon to the cell body.

**Figure 6.**
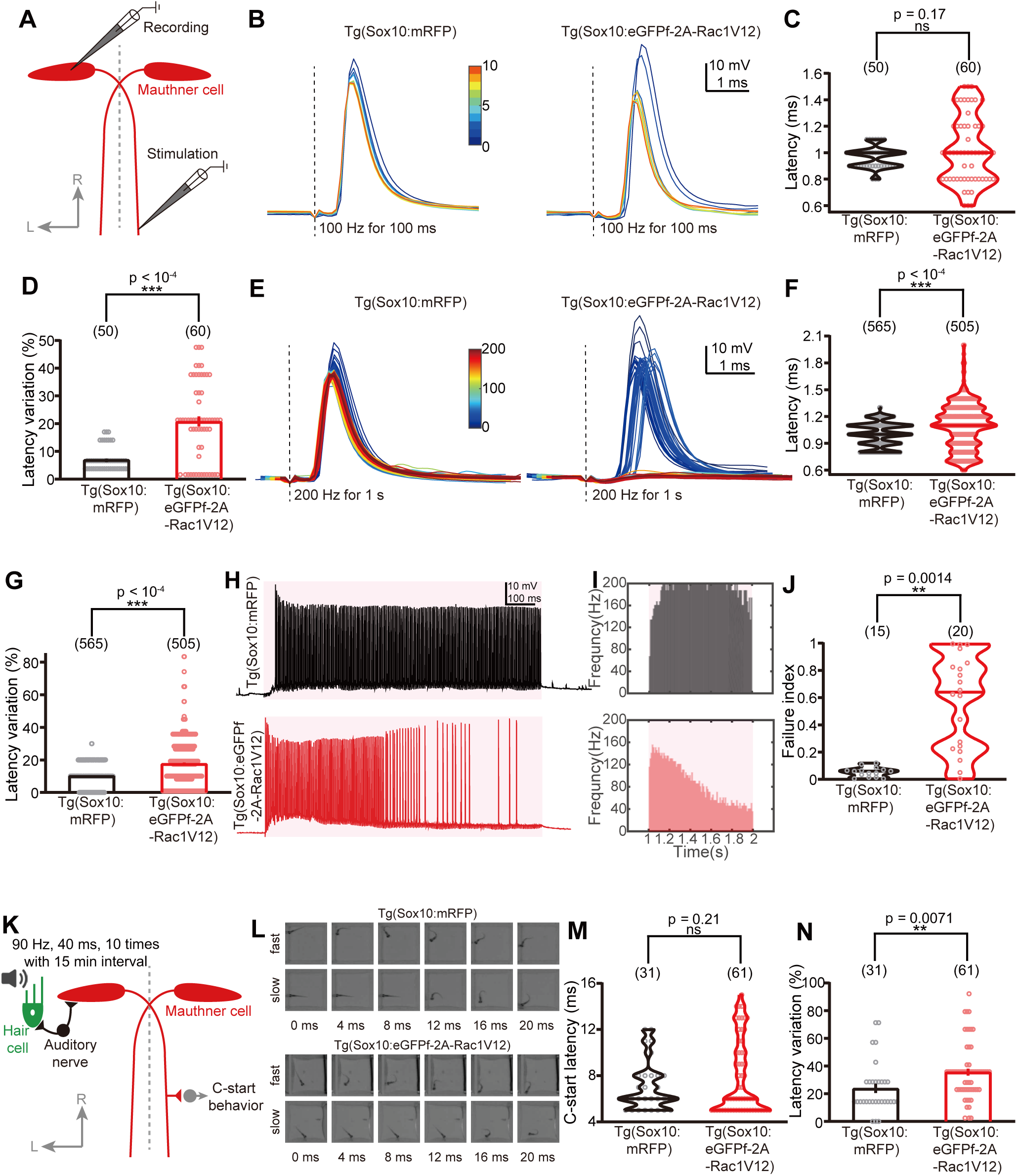
Disruption of anterior extension preference impairs proper axon functions. (A) Schematic of *in vivo* whole-cell recording of M-cells at 7 dpf. Rostral up. (B) Representatives of APs evoked by electrical stimulation (100 Hz, 100 ms) of M-axons in Tg(Sox10:mRFP) (left) and Tg(Sox10:eGFPf-2A-Rac1V12) (right) larvae. Dotted lines indicate the timing of electrical stimulation. Blue to red colors indicate the first to last evoked APs. (C) Latency of APs evoked by electrical stimulation (100 Hz, 100 ms) of M-axons in Tg(Sox10:mRFP) and Tg(Sox10:eGFPf-2A-Rac1V12) larvae. Data were obtained from 50 evoked-APs in 5 Tg(Sox10:mRFP) larvae and 60-evoked APs in 6 Tg(Sox10:eGFPf-2A-Rac1V12) larvae. The number on each column represents the number of evoked APs. (D) Latency variation of APs evoked by electrical stimulation (100 Hz, 100 ms) of M-axons in Tg(Sox10:mRFP) and Tg(Sox10:eGFPf-2A-Rac1V12) larvae. Same data set with (C) was used. (E) Representatives of APs evoked by HFS (200 Hz, 1 s) of M-axons in Tg(Sox10:mRFP) (left) and Tg(Sox10:eGFPf-2A-Rac1V12) (right) larvae. Dotted lines indicate the timing of electrical stimulation. Blue to red colors indicate the first to last evoked APs. (F) Latency of APs evoked by HFS (200 Hz, 1 s) of M-axons in Tg(Sox10:mRFP) and Tg(Sox10:eGFPf-2A-Rac1V12) larvae. Data were obtained from 565 evoked APs in 3 Tg(Sox10:mRFP) larvae and 505 evoked APs in 4 Tg(Sox10:eGFPf-2A-Rac1V12) larvae. The number on each column represents the number of evoked APs. (G) Latency variation of Aps evoked by HFS (200 Hz, 1 s) of M-axons in Tg(Sox10:mRFP) and Tg(Sox10:eGFPf-2A-Rac1V12) larvae. Same data set with (F) was used. (H) Samples of APs evoked by HFS (200 Hz, 1 s) of M-axons in Tg(Sox10:mRFP) (top) and Tg(Sox10:eGFPf-2A-Rac1V12) (bottom) larvae. Pink frame indicates the duration of HFS. (I) Average firing frequency (in 10-ms bins) of APs evoked by HFS (200 Hz, 1 s) of M-axons in Tg(Sox10:mRFP) (top) and Tg(Sox10:eGFPf-2A-Rac1V12) (bottom) larvae. Pink frame indicates the duration of HFS. Data were obtained from 3 Tg(Sox10:mRFP) and 4 Tg(Sox10:eGFPf-2A-Rac1V12) larvae. (J) Failure index of APs evoked by HFS (200 Hz, 1 s) of M-axons in Tg(Sox10:mRFP) and Tg(Sox10:eGFPf-2A-Rac1V12) larvae. Data were obtained from 15 trails in 3 Tg(Sox10:mRFP) embryos and 20 trails in 4 Tg(Sox10:eGFPf-2A-Rac1V12) embryos. The number on each column represents the number of trials used for statistics. (K) Schematic of sound-evoked C-start escape behavior at 7 dpf. Rostral up. (L) Representative images showing sound-evoked escape behavior of 7-dpf Tg(Sox10:mRFP) and Tg(Sox10:eGFPf-2A-Rac1V12) larvae in response to a sound stimulus, which was applied at 0 ms. (M) Latency of sound-evoked escape behavior of Tg(Sox10:mRFP) and Tg(Sox10:eGFPf-2A-Rac1V12) larvae at 7 dpf. Data were obtained from 31 trials in 13 Tg(Sox10:mRFP) embryos and 61 trails in 17 Tg(Sox10:eGFPf-2A-Rac1V12) embryos. The number on each column represents the trial number used for statistics. (N) Latency variation of sound-evoked escape behavior of Tg(Sox10:mRFP) and Tg(Sox10:eGFPf-2A-Rac1V12) larvae at 7 dpf. Same data set with (M) was used. Error bars, SEM. ns, not significant; **\***p < 0.05, **\*\***p < 0.01, **\*\*\***p < 0.001 (unpaired two-tailed Student’s t-test for (C), (D), (F), (G), (J), (M) and (N)). See also Figure S7.

We stimulated M-axon by 100 Hz for 100 ms and found no significant difference in the average latency of evoked APs (Figures 6B and 6C). However, the latency variation, which was calculated as the absolute value of the subtraction of single AP latency and average latency divided by the average latency, increased dramatically in Tg(Sox10:eGFPf-2A-Rac1V12) larvae (Figure 6D). Furthermore, by 100 Hz stimulation of M-axon for 2 s, the latency and latency variation of evoked APs both increased significantly in Tg(Sox10:eGFPf-2A-Rac1V12) larvae (Figures S7B-S7D). Furthermore, we monitored AP conduction under high-frequency stimulations (HFS) (200 Hz, 1 s) and found that APs could follow perfectly in control larvae, while in Tg(Sox10:eGFPf-2A-Rac1V12) animals, the firing frequency reduced significantly, especially in the last 400 ms (Figures 6E, 6H and 6I). The latency and latency variation of APs evoked by HFS were also increased notably (Figures 6F and 6G). Moreover, the failure index, which was calculated as the subtraction of total number of stimuli and the number of evoked APs divided by number of stimuli, increased markedly in Tg(Sox10:eGFPf-2A-Rac1V12) larvae (Figure 6J), suggesting that down-regulation of the anterior extension of myelin decreases faithful transduction of APs along M-axons.

As M-cells mediate C-start escape behavior of zebrafish, we next performed sound-evoked escape test at 7 dpf (Figure 6K). Although there was no significant difference in the average latency of C-start behavior (Figure 6L and 6M), the variation of C-start latency, which was calculated as the absolute value of the subtraction of single C-start latency and average latency divided by the average latency, increased markedly in Tg(Sox10:eGFPf-2A-Rac1V12) animals (Figure 6N). These results suggest that the defect of myelin anterior extension would reduce the reliability of C-start behavior.

## DISCUSSION

In the CNS, the conversion from oligodendroglial membranous sheets to well-established myelin segments is an extremely complex and highly regulated process^32,33^. According to the widely accepted model for the progression of myelination, after contact formation of an OL process with an axon, the growth pattern of developing myelin segments can be divided into two different motions: the spiral wrapping of the innermost layer of myelin sheath around axons and the bidirectional lateral extension along axons^34,35^. The spiral wrapping has been well elucidated to be driven by PI(3,4,5)P3-dependent polarized growth at the inner tongue^10^. However, few evidence has been provided to support the bidirectional lateral extension suggested by this model. Here, surprisingly, we found myelin sheaths exhibited anterior extension preference in zebrafish ventral SC as a novel process for precise regulation of myelination (Figure 7).

**Figure 7.**
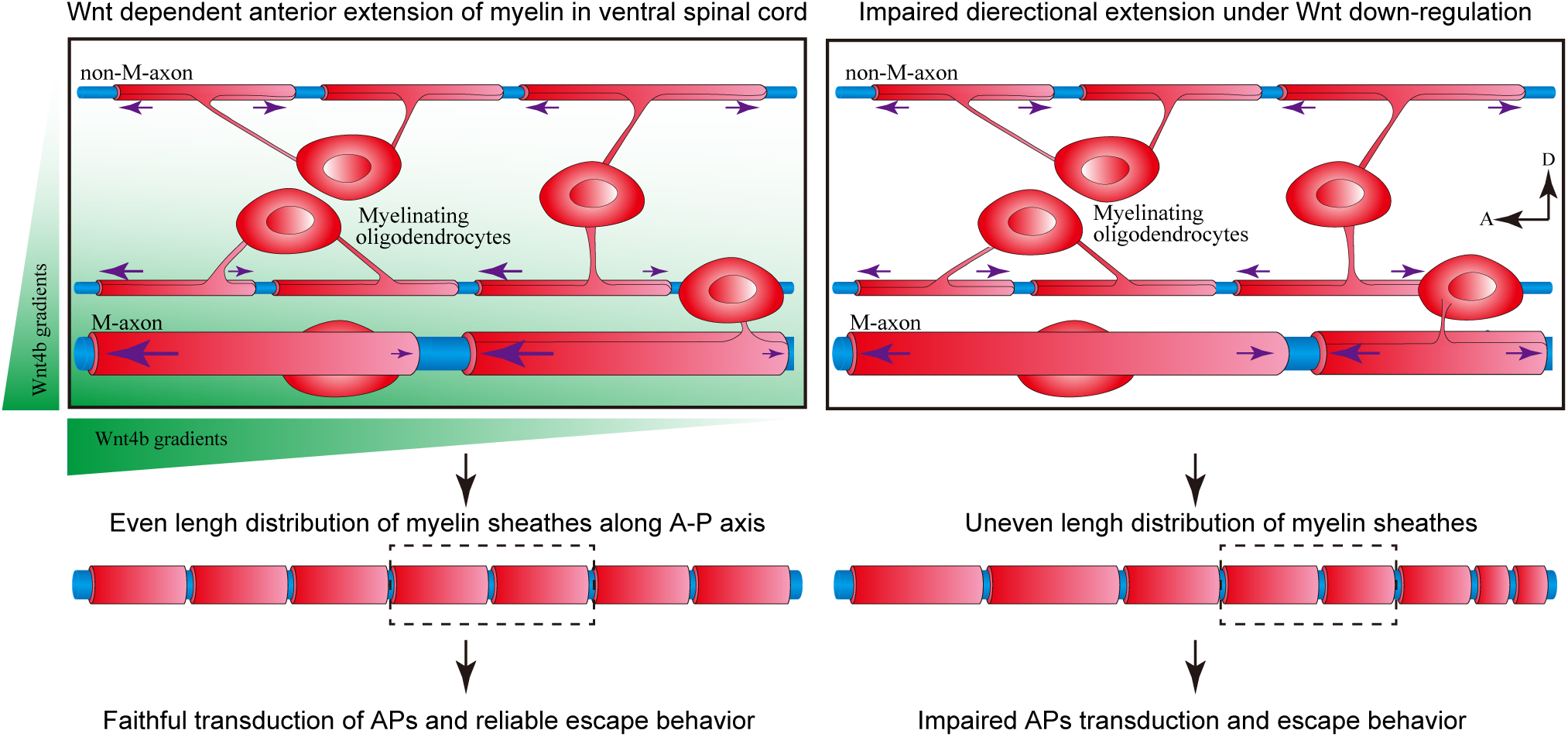
Working model. Anterior-to-posterior decreasing Wnt4b gradient leads to the anterior extension preference of myelin sheaths in the ventral SC, which contributes to the even length distribution of myelin segments along proximal-to-distal axis (left) and proper axon functions. Deficiency in myelin unidirectional extension by disruption of Wnt signaling impairs the uniformity of developing myelin sheaths, decreases faithful transduction of APs along M-axons and reduces the reliability of C-start behavior. The top panels are adopted from the dotted frames in the middle panels.

Wnt gradients have been well documented to contribute to the wiring establishment of neural circuitry in the CNS during development^36^. A decreasing anterior-to-posterior Wnt4b gradient was found to control the appropriate anterior turning of post-crossing commissural axons through an attractive mechanism via Frizzled3 in the ventral SC^24^. In addition, Wnt1 and Wnt5 gradients have been shown to be repulsive signals mediated by the Ryk receptor to promote the posterior-directed growth of descending corticospinal tract axons along the anterior-to-posterior axis in the SC of neonatal rodents and Drosophila^37–39^. Here, our study demonstrates that developing myelin sheaths exhibit anterior extension preference, which is dependent on an anterior-to-posterior Wnt4b gradient in the SC. These results indicate that Wnt gradient is not only essential for the neuronal wiring^24^, but also crucial for the sequential myelination along target axons, thereby facilitating the maturation, maintenance and plasticity of neural circuits (Figure 7).

OLs sampled M-axons and other small-calibre non-M-axon fibres during the first several hours after initiating myelination^40^. Anterior unidirectional extension of myelin sheaths is not specific to M-axons but a general phenomenon, as the non-M-axon-wrapping myelin in the ventral SC also exhibits similar growth preference. However, the growth PI of non-M-axon-wrapping myelin decreased significantly, which means that unlike the rigorous anterior extension of the myelin sheath along M-axons, the non-M-axon-wrapping myelin grows to both directions, with a preference to the anterior end. One possible mechanism underlying this difference is that the diameter of M-axons is up to ten times than that of non-M-axon fibres. As a result, the M-axon-wrapping myelin sheaths are more sensitive to the anterior-to-posterior Wnt4b gradient. Alternatively, our data do not exclude the possibility that Wnt-independent forces also contribute to the myelin directional extension. Previous studies have characterized many axonal cues, such as the polysialylated neural cell adhesion molecule^41^, the transmembrane protein Lingo1^42^ and neuregulins^43–46^, as instructive or inhibitory factors for myelination. Thus, it will be interesting to examine whether these axonal cues are required for myelin directional extension.

Indeed, it is interest to elucidate the underlying physiological function of myelin directional extension. In the vertebrate CNS, OL migration, maturation, and myelination occur along a proximal to distal gradient^40,47^. Therefore, for a newly formed myelin sheath, the already-formed myelin segment would be more likely to be present on the rostral side than on the caudal side along the M-axon. The rostral myelin would provide a stereospecific blockade to inhibit the anterior extension of this newly formed segment. Therefore, bidirectional extension would lead to much longer in length of myelin segments on the rostral side than on the caudal side, whereas the anterior extension would facilitate the even length distribution of myelin segments along axons. To test this hypothesis, we examined the length of myelin sheaths at the late stage of myelination, and found that in Tg(Sox10:eGFPf-2A-Rac1V12) transgenic line with impairment of growth preference, the length of myelin sheaths decreased significantly along proximal-to-distal axis, with longer in rostral side and shorter in caudal side. Thus, our findings imply that anterior extension preference contributes to the even length distribution of myelin sheaths along axons during the developmental myelinating stage.

In vertebrates, the saltatory conduction of AP along myelinated axons requires appropriate length of myelin segments. Previous studies have showed that the conducting velocity of AP follows an inverse-U-shaped curve related to myelin length^48,49^. A mathematical spike-diffuse-spike modelling of AP propagation demonstrated that the optimal internode length ensures the optimal AP conduction^48^. Moreover, a previous study demonstrated that too long or too short myelin length could both impair the conducting velocity of AP^49^. Therefore, the uniformity to optimal length of myelin segments along proximal-to-distal axis contributes to the appropriate saltatory conduction of AP. In this study, we found that in Tg(Sox10:eGFPf-2A-Rac1V12) transgenic line with impairment of even length distribution of myelin sheaths, the faithful transduction of AP on M-axon and the reliability of escape behavior were both reduced significantly. In the behavioral test, although the mean latency of escape responses remained unchanged, the increased variation in latency indicates that, when facing a predator, the animal’s escape speed may, with a certain probability, be markedly slower than the average, which is critically deleterious to animal survival. Our study offers a mechanism underlying the formation of myelin length uniformity and shows that deficiencies in myelin length uniformity compromise the proper axon functions.

In summary, our data show that Wnt-gradient-mediated myelin preferentially unidirectional extension acts as a novel process for precise regulation of myelination during development, providing new insight into myelin structuring and functioning.

## RESOURCE AVAILABILITY

### Lead contact

Further information and requests for resources and reagents should be directed to and will be fulfilled by the lead contact, Cheng He.

### Materials availability

Plasmids and fish lines generated in this study are available from the lead contact without restriction.

### Data and code availability

This paper does not report any original code.

Any additional information required to reanalyze the data reported in this work paper is available from the lead contact upon request.

## ACKNOWLEDGEMENTS

We are grateful to Dr. Bruce Appel for providing the plasmid *Sox10:mRFP*, Dr. Koichi Kawakami for providing the Tg(hspGFF62A,UAS:GFP) line, Dr. Zilong Weng for providing the Tol056 transgenic line, and Dr. Xu Wang for providing the Tg(Hsp70:Dkk1b-GFP) transgenic line.

## Funding

Ministry of Science and Technology of the People’s Republic of China STI2030-Major Projects 2022ZD0204700 (to C.H.)

National Natural Science Foundation of China 31400920 (to P.L.)

National Natural Science Foundation of China 31871026 (to C.H.)

National Key Research and Development Program of China 2016YFA0100802 (to C.H.)

Shanghai Municipal Science and Technology Major Project 18JC1410100 and 2018SHZDZX05 (to J.L.D.)

Key Research Program of Frontier Sciences of Chinese Academy of Sciences QYZDY-SSW-SMC028 (to J.L.D.)

Strategic Priority Research Program of Chinese Academy of Sciences XDB32010200 (to J.L.D.)

## AUTHOR CONTRIBUTIONS

P.L., J.L.D. and C.H. designed the experiments and wrote the manuscript. J.L.D. and C.H. conceived and supervised the research project. P.L., Q.S.D., Q.C. and X.C. performed the experiments and analyzed the data.

## STAR★METHODS

Detailed methods are provided in the online version of this paper and include the following:

- EXPERIMENTAL MODEL AND SUBJECT DETAILS

- Zebrafish
- METHOD DETAILS

- In Situ Hybridization
- DNA Microinjection
- Plasmid Construction and Generation of Transgenic Zebrafish Lines
- Morpholino Oligomer-mediated Knockdown
- Time-lapse Confocal Imaging and Analysis
- Photo-Conversion and Imaging
- Infrared Laser Local Heat-shock Irradiation
- Transmission Electron Microscopy
- Drug Administration
- In Vivo Electrophysiological Recording
- Escape Behavior Test
- QUANTIFICATION AND STATISTICAL ANALYSIS

- Statistics

## SUPPLEMENTAL INFORMATION

Supplemental Information includes Figures S1-S7.

## STAR★METHODS

### EXPERIMENTAL MODEL AND SUBJECT DETAILS

#### Zebrafish

Adult zebrafish (*Danio rerio*) were maintained at the National Zebrafish Resources of China (NZRC, Shanghai, China) with an automatic fish housing system (ESEN, Beijing, China) and following standard protocols^50,51^. Zebrafish embryos were raised at 28°C and staged according to hours (or days) post-fertilization or morphological criteria. Embryos and larvae were raised under a 14 hr - 10 hr light-dark cycle in 10% Hank’s solution, which consisted of (in mM) 140 NaCl, 5.4 KCl, 0.25 Na_2_HPO_4_, 0.44 KH_2_PO_4_, 1.3 CaCl_2_, 1.0 MgSO_4_ and 4.2 NaHCO_3_ (pH 7.2), and were treated with 0.003% 1-phenyl-2-thiourea (PTU, Sigma) to prevent pigment formation. The handling procedures were approved by the Institute of Neuroscience, Chinese Academy of Sciences. The transgenic zebrafish strains Tg(Olig2:eGFP), Tg(Olig2:Kaede), Tg(Sox10:mRFP), Tg(Flk1:eGFP), Tg(Sox10:eGFPf-2A-Rac1V12), Tg(Sox10:mRFP-2A-Lrp6dC), Tg(hspGFF62A,UAS:GFP)^52^, Tol056^53^, Tg(Hsp70:Dkk1b-GFP)^54^ and Tg(Hsp70:Wnt4b-2A-eGFP) were used in this study.

Knockout zebrafish were constructed through CRISPR-Cas9 technology, and the following sgRNA sequence were used: Wnt4b sgRNA: CACGTTGATCCCCCGAGGTGTGG; Lrp6 sgRNA: GCGTCAGGCCGAACGGGTGCGGG. Cas9 protein (NEB, M0646M) and the sgRNA were microinjected into the animal pole of one-cell stage fish embryos. Injected F0 animals were raised to adulthood and outcrossed to wild-type animals to create F1 offspring. Clutches of F1 offspring were raised to adulthood and genotyped to identify heterozygous carriers of function disrupting mutant alleles (Figure S5).

### METHOD DETAILS

#### *In Situ* Hybridization

To synthesize antisense *Wnt4b* probes, the entire coding region of *Wnt4b* was isolated by PCR from whole-embryo cDNA generated at 2 dpf by using an oligonucleotide pair with the following sequences^25^: forward, *5’-ATGCCAACAGTCTCCTCTGTGACTC-3’*; reverse, *5’-TTATTCTCGGCAGGTGTGTATGAGC-3’*. Then, the fragment was subcloned into *pGEM-T* easy vector (Promega) followed by in vitro transcription by using the digoxigenin (DIG) RNA labelling kit (Roche). Whole-mount *in situ* hybridization was performed as described previously^55^. Embryos at 1 - 4 dpf were treated with proteinase K (10 μg/ml) for 15, 30, 45, and 60 min, respectively. *Wnt4b* probe at 3 ng/μl was hybridized to embryos in 150 μl of hybridization buffer for 12.5 hr at 65 °C. The anti-DIG antibody incubation was performed for 15 hr at 4 °C. Embryos were mounted in glycerol, and high-magnification pictures were taken using a microscope (Nikon E600FN) with a digital camera.

The transversal histological analysis of the SC was conducted based on vibratome sections. The larval samples by whole-mount *in situ* hybridization were embedded in 4% low-melting point agarose and 100 μm-thick sections were cut using Leica VT 1200S. Images were obtained on an Olympus microscope (1X71) mounted with a DP72 digital camera controlled by DP2-BSW software. For quantification of the *Wnt4b* signal intensity in transversal section, the images were taken at the same exposure and converted into grey scale. The ventral SC areas were circled by a threshold eliminating background values and mean grey values were obtained. The intensity indicated by the mean grey values in the ventral SC area represents the relative expression levels. The Y-axis of the right panel in Figure 2H indicates the relative *Wnt4b* expression level of each section from the left panel, normalized to the baseline level (section 1).

#### DNA Microinjection

Fertilized eggs were microinjected at the one-cell stage with 1 nl of solution containing 25 ng/μl *Sox10:mRFP*, *Sox10:eGFP*, *Sox10:Kaede*, *Sox10:mRFP-2A-Lrp6dC*, *Sox10:eGFPf-2A-Rac1V12*, or *Sox10:eGFPf-2A-hCRIB-RasCT* plasmid DNA and I-SceI (New England Biolabs) as described^56^. The *Tol2-Hsp70:Wnt4b-2A-eGFP* plasmid was injected at a concentration of 20 ng/μl in combination with *Tol2* transposase mRNA at 25 ng/μl into one-cell embryos^57^.

#### Plasmid Construction and Generation of Transgenic Zebrafish Lines

To create the plasmid *Sox10:eGFP*, the coding sequence of *eGFP* was PCR amplified from *GFAP-eGFPf* and then sub-cloned into *Sox10:mRFP-pBSII sk(+)* to replace *mRFP*. To create the plasmid *Sox10:Kaede,* the coding sequence of *kaede* was PCR amplified from *pCS2-kaede* and the farnesyl fragment was PCR amplified from *GFAP-eGFPf*. The two fragments were then sub-cloned into *Sox10:mRFP-pBSII sk(+)* to substitute *mRFP*. The plasmids *Sox10:mRFP-2A-Lrp6dC* were generated as following: A *mRFP* fragment was PCR amplified from *Sox10:mRFP* and the *Lrp6dC* was PCR amplified from *Lrp6dC-PCS*^23^. The *mRFP* and *Lrp6C* fragments were then sub-cloned into *Sox10:mRFP-pBSII sk(+)* to replace *mRFP* and generate the required plasmid *Sox10:mRFP-2A-Lrp6dC*. To create the plasmid *Tol2-Hsp70:Wnt4b-2A-eGFP*, the entire coding region of *Wnt4b* was PCR amplified from *Wnt4b-pGEMT-easy* and the *eGFP* fragment was PCR amplified from *pCS2-eGFP*. The two fragments were then sub-cloned into *Tol2-Hsp70:mRFP-Cre* to replace *mRFP-Cre* and generate the required *Tol2-Hsp70:Wnt4b-2A-eGFP* plasmid. To construct the plasmids *Sox10:eGFPf-2A-hCRIB-RasCT* and *Sox10:eGFPf-2A-Rac1V12*, an *eGFPf* fragment was PCR amplified from *GFAP-eGFPf* and the *hCRIB-RasCT* (and *Rac1V12*) fragment was PCR amplified from *hCRIB-RasCT-nos1-3’UTR* (and *Rac1V12-nos1-3’UTR*)^30^. The two fragments were then sub-cloned into *Sox10:mRFP* to replace *mRFP* and generate the required plasmids. The transgenic line Tg(Hsp70:Wnt4b-2A-eGFP) was created by injecting *tol2* DNA plasmid *Tol2-Hsp70:Wnt4b-2A-eGFP*, together with *tol2* mRNA, into one-cell embryos. The transgenic lines Tg(Sox10:mRFP), Tg(Sox10:eGFPf-2A-Rac1V12), Tg(Sox10:mRFP-2A-Lrp6dC), was created by injecting plasmid *Sox10:mRFP*, *Sox10:eGFPf-2A-Rac1V12*, *Sox10:mRFP-2A-Lrp6dC*, together with I-SceI enzyme, into one-cell embryos.

#### Morpholino Oligomer-Mediated Knockdown

The lyophilized MOs were diluted in nuclease-free water (Ambion) to 1 mM, and concentrations were checked by spectrophotometry (A265 in 0.1 N HCl) according to the standard protocol of Gene Tools. Stocks were then diluted to various working concentrations with Danieau’s buffer (58 mM NaCl, 0.7 mM KCl, 0.4 mM MgSO_4_, 0.6 mM Ca(NO_3_)_2_, 0.5 mM HEPES, pH 7.6), and 1 nl was pressure-injected into one-cell-stage embryos using a Picospritzer II micro-injector. MOs sequences were as follows:

Control MO: *5’-CCTCTTACCTCAGTTACAATTTATA-3’*

*β-catenin* MO^58^: *5’-ATCAAGTCAGACTGGGTAGCCATGA-3’*

*Wnt1* MO^59^: *5’-AGCAACGCGAGAACCCGCATGATAT-3’*

*Wnt3* MO^60^: *5’-AATCCAACCAGGTACAAATCCATGA-3’*

*Wnt5* MO^61^: *5’-GTCCTTGGTTCATTCTCACATCCAT-3’*

*Wnt8 MO*^62^*: 5’-ACGCAAAAATCTGGCAAGGGTTCAT-3’*

*Wnt11* MO^63^: *5’-GAAAGTTCCTGTATTCTGTCATGTC-3’*

*Wnt4b* MO^64^: *5’-GTTGGCATCAGATTGCCTGTCTGTC-3’*

#### Time-Lapse Confocal Imaging and Analysis

Embryonic and larval zebrafish during imaging were anesthetized using 3-aminobenzoic acid ethyl ester, embedded in 1% low-melting-point agarose (Sigma), and then mounted on their sides in glass-bottomed 35-mm Petri dishes^65,66^. Immediately after imaging at each time point, the fish were removed to rearing solution for recovery. Time-lapse images were captured using 40X water-immersion (NA: 0.80) objectives mounted on an Olympus FV1000-MPE laser-scanning confocal microscope (Tokyo, Japan) equipped with a heated chamber to maintain the embryos at 28°C. Z-stack images were collected with an interval of 12 hr. Image-Pro Plus (Media Cybernetics, Bethesda, MD) and ImageJ (NIH) were used to analyse the time-lapse imaging data. The analyses of myelin segments were carried out on Z-stack images by using unbiased stereological methods. Early-stage myelin segments were defined as discrete rod-like expansions along OL processes. The Z-stack images were collected in 1 μm z-steps. The anterior and posterior ends of myelin segments at each time-point were confirmed by slice-to-slice characterization of Z-stack images. Black background was used for the alignment of time-lapse images. Multiple images were partially overlapped to obtain a complete myelin segment at late developmental stage. We used Growth PI, which was calculated as the subtraction of extension length in anterior and posterior ends divided by total extension length as follows, to characterize the directional extension preference of myelin sheaths.

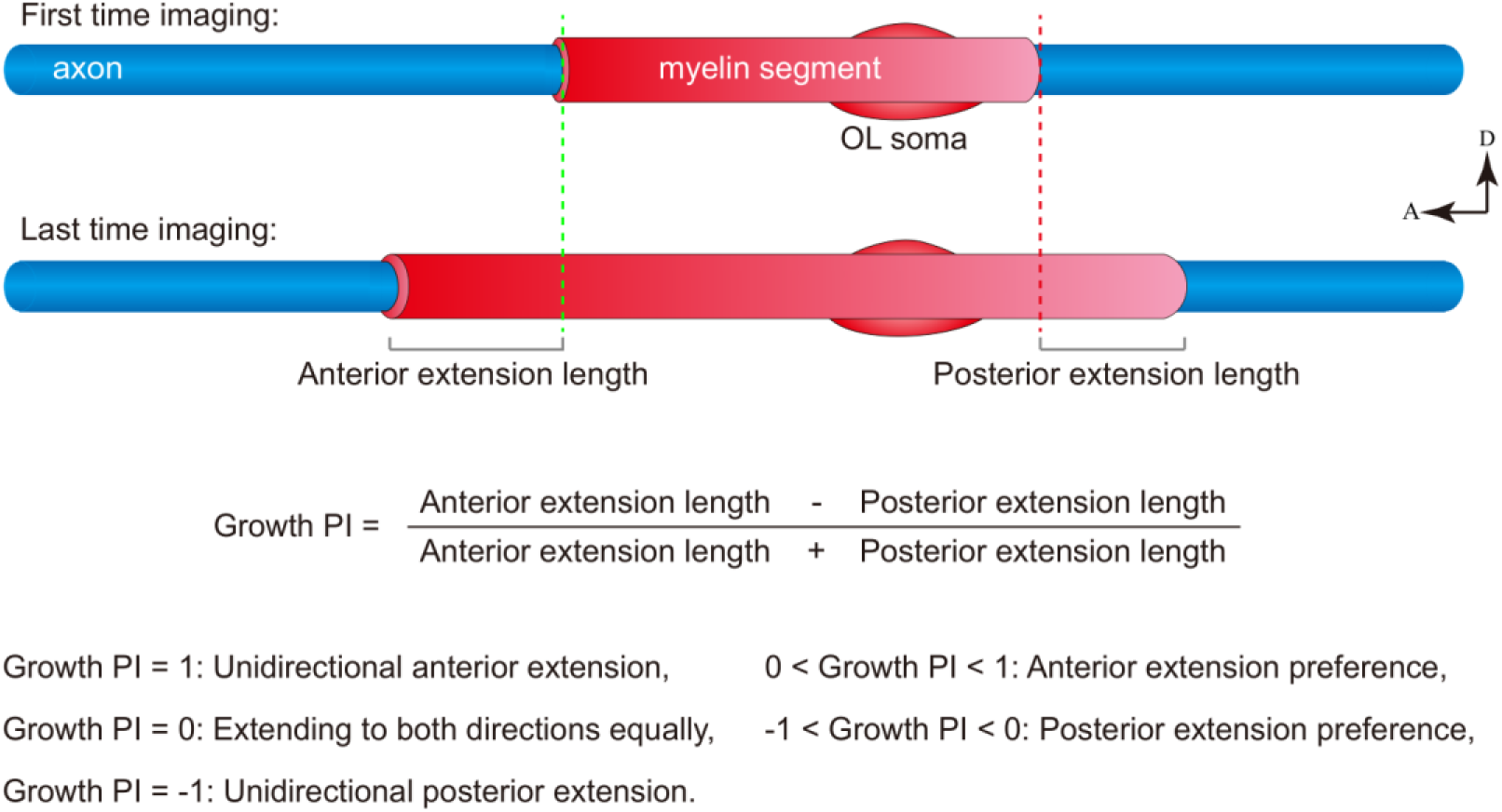

#### Photo-Conversion and Imaging

All photoconversion experiments and fluorescent imaging were performed using Olympus FV1000-MPE (Tokyo, Japan) confocal microscope. To block pigmentation when imaging embryos older than 24 hpf, embryos were raised in the presence of 0.003% PTU from approximately 22 hpf. For photo-conversion and imaging, embryos were mounted in 0.5% low melting point agarose (Lonza, 50101) in 30% Danieau’s solution. Embryos were imaged before photo-conversion using a 488 nm laser using the minim90al necessary laser power. Photo-conversion of Kaede was performed by scanning the selected region of interest with a 405 nm laser. The scans were repeated until green fluorescence was eliminated and red fluorescence was confirmed and documented. Embryos were then released from the agarose and transferred to a petri dish with egg water containing PTU and covered with aluminum foil to protect from light. Embryos were kept at 28°C until they were imaged again.

#### Infrared Laser Local Heat-Shock Irradiation

The optical system used for the infrared laser local heat-shock irradiation was described in a previous report^27^. An infrared optical unit and a mono-coated objective lens were used for irradiation. The power of the infrared laser was measured before irradiation. During the trials of infrared laser irradiation of the targets, the Tg(Hsp70:Dkk1b-GFP) and Tg(Hsp70:Wnt4b-2A-eGFP) larvae were embedded in a 3% methylcellulose solution for stable positioning of the targets and anesthetized with 0.02% MS222 (M0387; Sigma). To prevent pigmentation by melanin synthesis, the zebrafish embryos were grown in 0.003% PTU. Infrared laser irradiation was carried out under observation with a microscope to distinguish the to-be-targeted zebrafish larvae somite. Laser irradiated for 10 s each time. The green fluorescence after every irradiation was checked when GFP began to express significantly. Enhance the fluorescence every day during the period of time-lapse imaging. After infrared irradiation, zebrafish larvae were kept under normal culture conditions before observation. The local infrared heat application was performed after the target oligodendrocyte had differentiated and begun to form myelin segments along M-axons, indicating that the reduction of myelin Growth PI by the manipulation of Wnt gradient was not mediated by oligodendrocyte differentiation.

#### Transmission Electron Microscopy

The spinal cord (SC) of zebrafish at 7 dpf were perfused with a phosphate buffer solution containing 2.5% glutaraldehyde and 4% PFA (pH 7.3). The samples were fixed in 1% osmium tetroxide for 45 min, dehydrated, embedded in Araldite resin. Then, the samples were sectioned to a thickness of 70-90 nm using a diamond knife (Leica EM UC7) to 70-90nm, followed by stained with uranyl acetate and lead citrate for contrast enhancement. All the samples were examined using a transmission electron microscope (HITACHI H-7650 or FEI Talos 120) at an accelerating voltage of 100 kV.

#### Drug Administration

The larvae were randomly divided into control and the antagonists-treated groups. They were then transferred into small petri dishes containing with SU5402 (Selleck, S7667, 10 µm), Cyclopamine (Selleck, S1146, 10 µm), LDN193189 (Selleck, S2618, 5 µm), PNU74654 (Selleck, S8429, 10 µm), NSC23766 (Calbiochem, 553502, 150 μM) or vehicle^67–69^. The solutions were renewed every day during the imaging period.

#### In Vivo Electrophysiological Recording

*In vivo* electrophysiological recording was performed as previously described^70^. Larvae were first paralyzed with the neuromuscular junction blocker α-bungarotoxin (1 mg/ml, Tocris) for 40 s - 1 min, and then embedded in 1.5% - 1.7% low melting-point agarose (Sigma-Aldrich). The extracellular solution consisted of (in mM): 134 NaCl, 2.9 KCl, 2.1 CaCl_2_, 1.2 MgCl_2_, 10 HEPES and 10 glucose (pH = 7.8). The internal solution consisted of (in mM): 100 K-Gluconate, 10 KCl, 2 CaCl_2_·2H_2_O, 2 Mg_2_·ATP, 0.3 GTP·Na_4_, 2 Phosphocreatine, 10 EGTA, 10 HEPES (pH 7.4). The equilibrium potential of chloride ions (E_Cl_^-^) was about −60 mV according to the Nernst equation. Recording micropipettes were made from borosilicate glass capillaries (BF100-58-10, Sutter Instrument). In vivo whole-cell recording of M-cells were made under visual guidance with a patch-clamp amplifier (MultiClamp 700B, Axon Instruments). Signals were filtered at 2 - 5 kHz and sampled at 10 kHz by using Clampex 10.2 (Molecular Devices). Electrical stimuli used for stimulating M-axons were 20 mV (0.1 ms in duration), delivered through a theta-micropipette (2 μm in tip diameter).

For whole-cell recording of M-cells, a tiny cut breaking the skin was made dorsally ∼ 100 μm rostral to the location of M-cells. A recording micropipette (6 - 12 MΩ in resistance, 1.5 - 2 μm in tip diameter) filled with the internal solution was inserted into the brain through this cut, and approached the M-cells from a caudal-ventral direction with a persistent positive pressure for keeping the tip clean. After the contact between the micropipette tip and M-cell soma, a giga-Ohm seal was formed by removing the positive pressure and applying a slight negative pressure. Whole-cell recording was then achieved by delivering a few brief electrical zaps (25 - 50 μs in duration) to break the cell membrane beneath the micropipette tip. In voltage clamp (v.c.) mode, M-cells were usually held at E_Cl_- (−60 mV).

#### Escape Behavior Test

Sound stimuli (90 dB, 40 ms) were delivered through a speaker with a plexiglass plate on it and one 96-well plate with one larva in each well was placed on the plexiglass plate. In most of experiments, the 96 wells were not fully loaded because the well wall in the corner prevents a clear view of the larvae. Larval behavior was monitored by using an infrared-sensitive high-speed camera at 1000 Hz (Zebrabox, Viewpoint). Larvae were allowed to freely swim in the test arena for 40 - 50 min to adapt before the first test was performed. During testing, the larval behavior was simultaneously recorded during an experiment. 10 trials in each test were carried out to calculate the latency of escape behavior and the trial-to-trial interval was 15 min. Successful C-start behavior was visually identified for each larva, and corresponded to the C-shape rapid movement within 5 – 20 ms after the sound onset^70^.

### QUANTIFICATION AND STATISTICAL ANALYSIS

#### Statistics

Statistical analysis was performed by using unpaired two-tailed Student’s *t*-tests, one-way ANOVA and two-way ANOVA test. P values less than 0.05 were considered to be statistically significant. All results are presented as the mean ± SEM.

---

**Figure S1.**
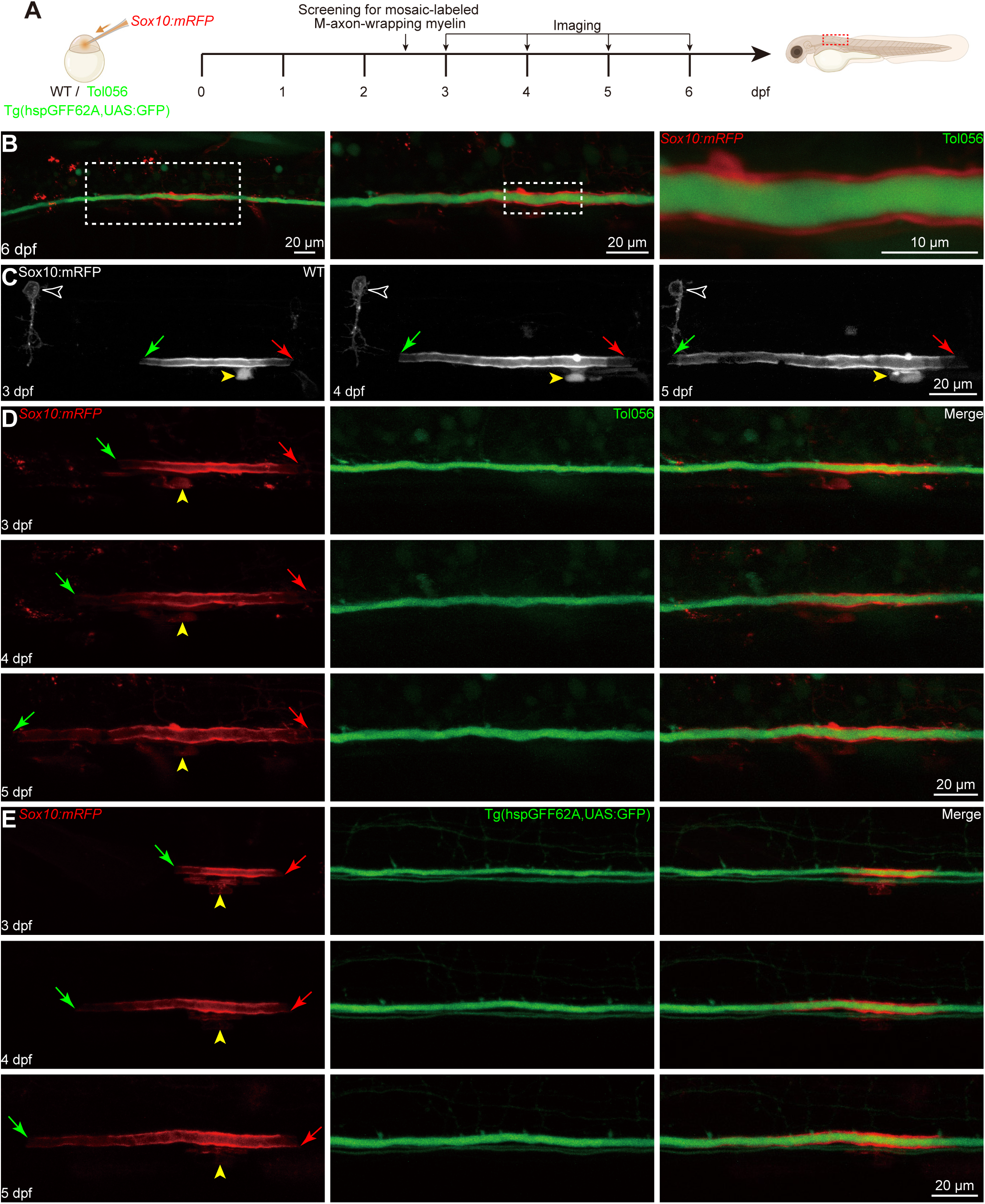
Anterior extension preference of M-axon-wrapping myelin in Tol056 and Tg(hspGFF62A,UAS:GFP) larvae. (A) Diagram of single M-axon-wrapping myelin sheath mosaically labeled by Sox10:eGFP in WT, Tol056 and Tg(hspGFF62A,UAS:GFP) lines. Timeline indicating the screening and imaging procedures. (B) Visualization of both myelin sheaths and M-axons. Left, projected image adopted from red dotted frame in the right cartoon in (A); Middle, confocal image enlarged from the boxed area in the left panel; Right, confocal image enlarged from the boxed area in the middle panel. (C) Time-lapse images of an individual OL with a M-axon-wrapping myelin segment. White hollow arrowhead, non-specific labelled neuron as reference marker; (D) Time-lapse images of M-axon-wrapping myelin in (B) in Tol056 larvae. Left, M-axon-wrapping myelin acquired from *Sox10:mRFP* larvae; Middle, M-axon in Tol056 larvae; Right, merged images of M-axon and myelin segments. (E) Time-lapse images of M-axon-wrapping myelin in Figure 1B in Tg(hspGFF62A,UAS:GFP) larvae. Left, M-axon-wrapping myelin acquired from *Sox10:mRFP* larvae; Middle, M-axon in Tg(hspGFF62A,UAS:GFP) larvae; Right, merged images of M-axon and myelin segments.

**Figure S2.**
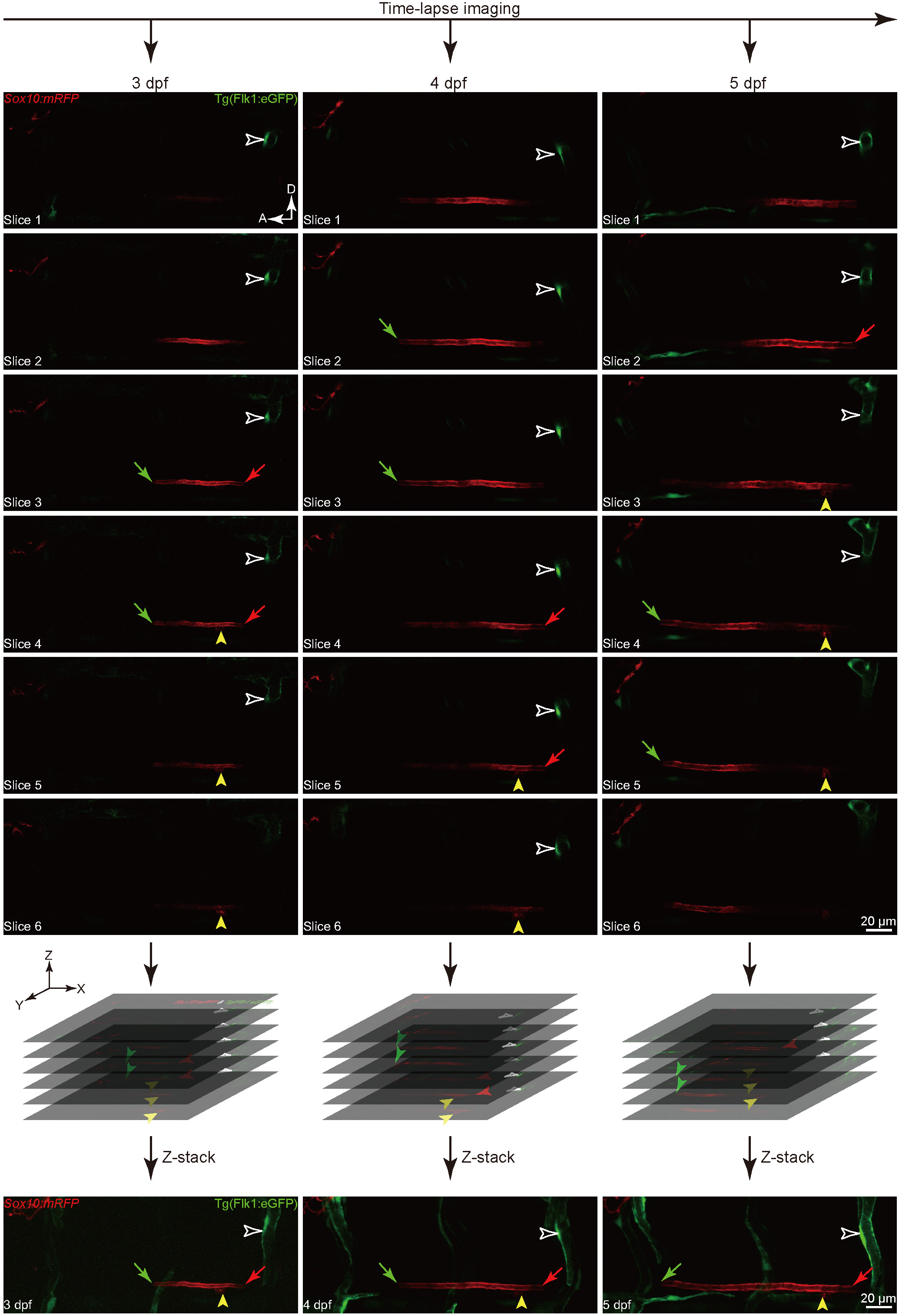
Confirmation of anterior and posterior ends of M-axon-wrapping myelin in the SC of Tg(Flk1:eGFP) larvae. A series of images of an individual M-axon-wrapping myelin at 3 (left), 4 (middle) and 5 (right) dpf. The Z-stack images were obtained using an Olympus FluoView 1000 laser-scanning confocal microscope, and images were collected in 1 μm z-step. The anterior (green arrows) and posterior (red arrows) ends of myelin segments at each time-point were confirmed by slice-to-slice characterization of Z-stack images. D, dorsal; A, anterior; White hollow arrowhead, ISVs as reference markers; Yellow arrowhead, OL soma as fixed point of reference markers during development.

**Figure S3.**
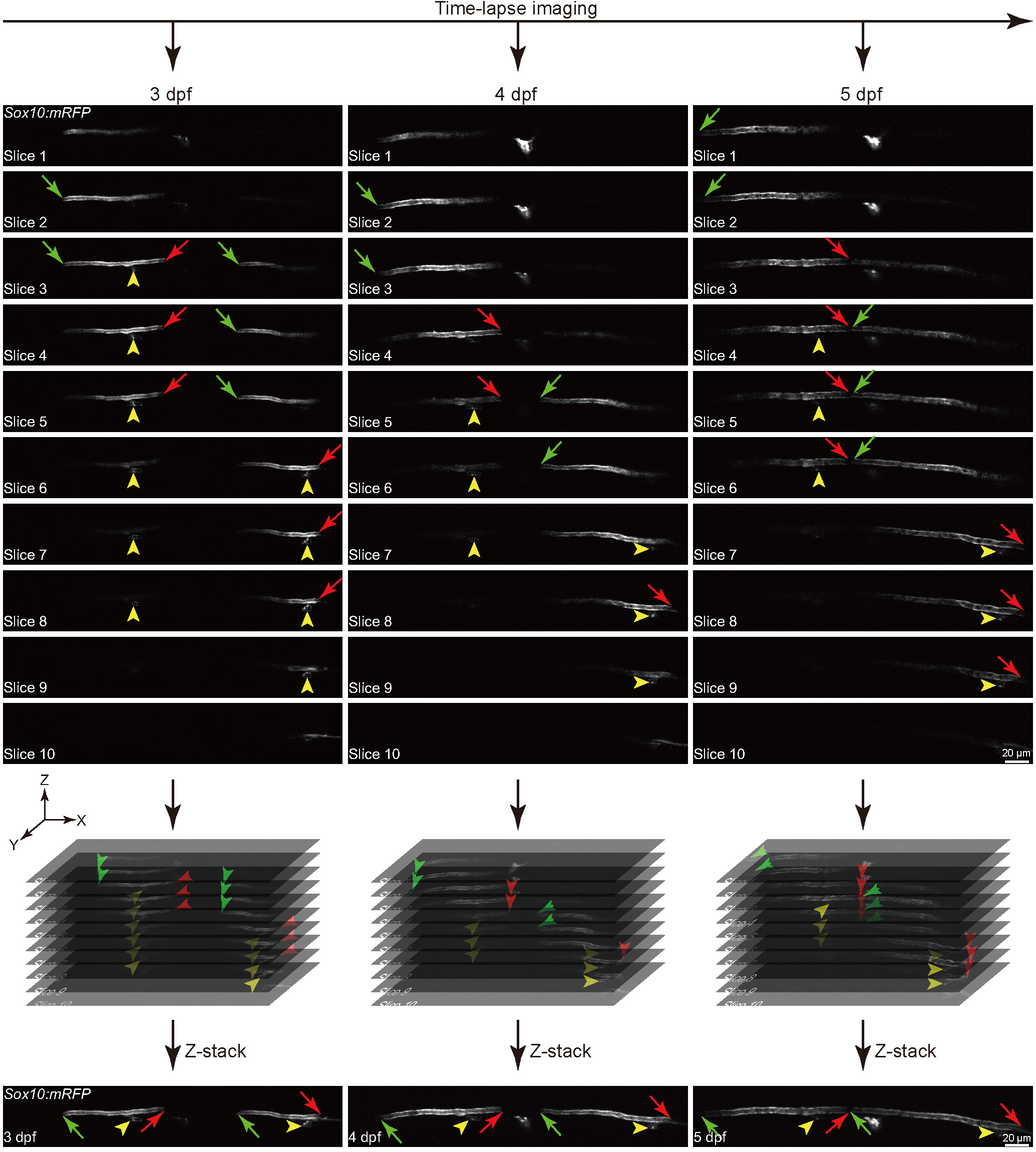
Confirmation of anterior and posterior ends of two neighboring myelin segments along the M-axon. A series of images of two neighboring M-axon-wrapping myelin segments at 3 (left), 4 (middle) and 5 (right) dpf. The Z-stack images were obtained using an Olympus FluoView 1000 laser-scanning confocal microscope, and images were collected in 1 μm z-step. The anterior (green arrows) and posterior (red arrows) ends of myelin segments at each time-point were confirmed by slice-to-slice characterization of Z-stack images. Yellow arrowhead, OL soma as fixed point of reference markers during development.

**Figure S4.**
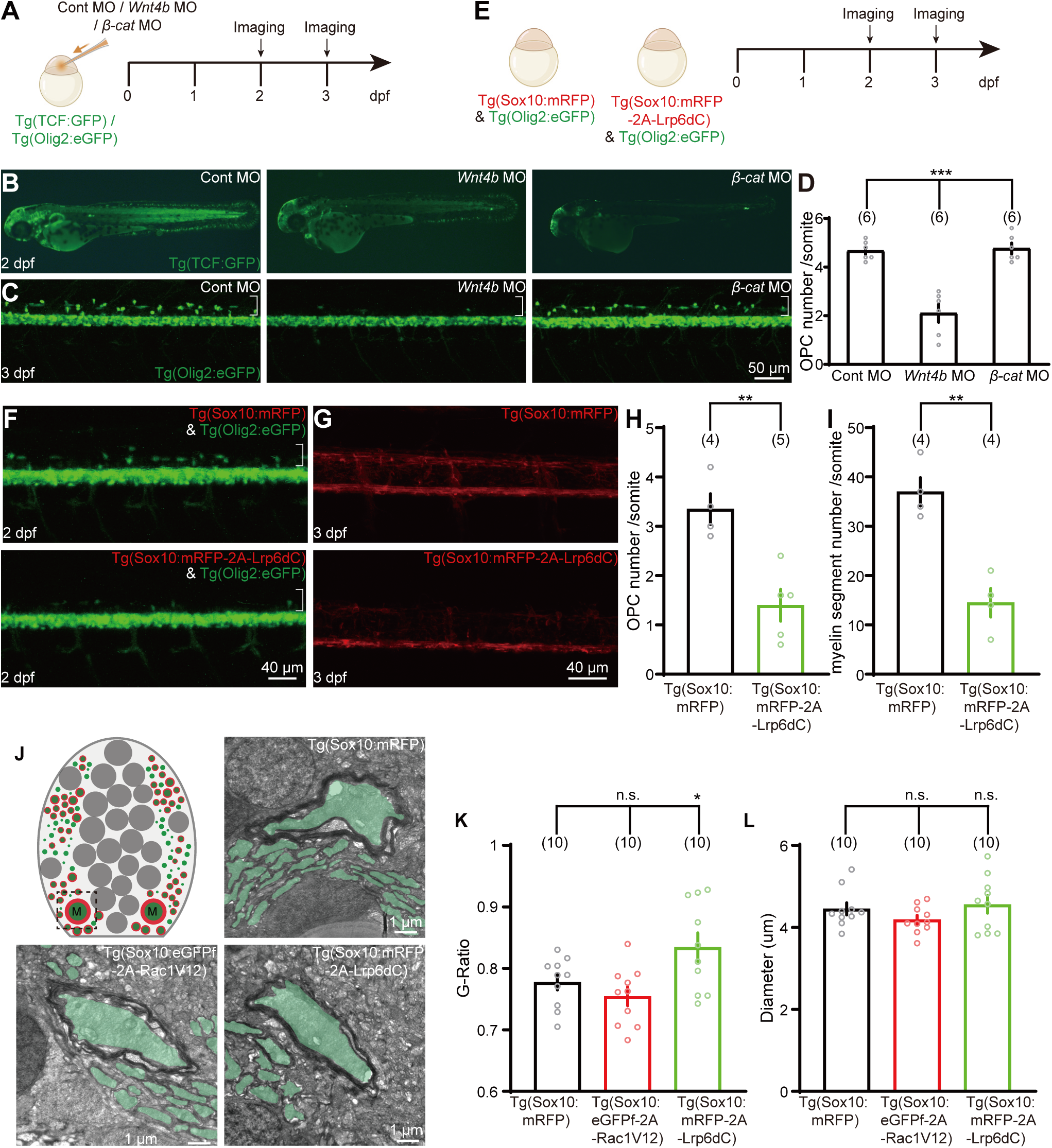
Requirement of Wnt signaling for oligodendroglial development and myelination. (A) Schematic diagram displaying the imaging timeline of Wnt activation and oligodendrocytes dorsal migration in control, *Wnt4b* and *β-catenin* morphants. (B) Confocal imaging of Wnt reporter line Tg(TCF:GFP) in control, *Wnt4b* and *β-catenin* morphants. (C) Confocal imaging of oligodendrocytes dorsal migration labelled by Tg(Olig2:eGFP) lines in control, *Wnt4b* and *β-catenin* morphants. The white bracket indicates the area of the dorsal SC. (D) OPC number per somite in the dorsal SC of Tg(Olig2:eGFP) lines in control, *Wnt4b* and *β-catenin* morphants. (E) Schematic diagram displaying the imaging timeline of oligodendrocytes dorsal migration in Tg(Sox10:mRFP) and Tg(Sox10:mRFP-2A-Lrp6dC) transgenic lines. (F) Confocal imaging of oligodendrocytes dorsal migration labelled by Tg(Olig2:eGFP) in Tg(Sox10:mRFP) and Tg(Sox10:mRFP-2A-Lrp6dC) transgenic lines. The white bracket indicates the area of the dorsal SC. (G) Confocal imaging of myelin segments in Tg(Sox10:mRFP) and Tg(Sox10:mRFP-2A-Lrp6dC) transgenic lines. (H) OPC number per somite in the dorsal SC of Tg(Sox10:mRFP) and Tg(Sox10:mRFP-2A-Lrp6dC) transgenic lines. (I) Myelin segments number per somite in the dorsal SC of Tg(Sox10:mRFP) and Tg(Sox10:mRFP-2A-Lrp6dC) transgenic lines. (J) Schematic and TEM images of the myelinated tracts in the ventral SC of Tg(Sox10:mRFP), T g(Sox10:mRFP-2A-Lrp6dC) and Tg(Sox10:eGFPf-2A-Rac1V12) larvae. Green area indicates myelinated axon. Dotted frame in Schematic indicates the imaging area. M, M-axons. (K) G-Ratio of M-axons of Tg(Sox10:mRFP), Tg(Sox10:mRFP-2A-Lrp6dC) and Tg(Sox10:eGFPf-2A-Rac1V12) larvae. (L) Diameter of M-axons of Tg(Sox10:mRFP), Tg(Sox10:mRFP-2A-Lrp6dC) and Tg(Sox10:eGFPf-2A-Rac1V12) larvae. The number on each column in (D, H, I, K and L) represents the number of embryos used for statistics. Error bars, SEM. ns, not significant; *p < 0.05, **p < 0.01, ***p < 0.001 (two-way ANOVA for (D, K and L); unpaired two-tailed Student’s t-test for (H and I)).

**Figure S5.**
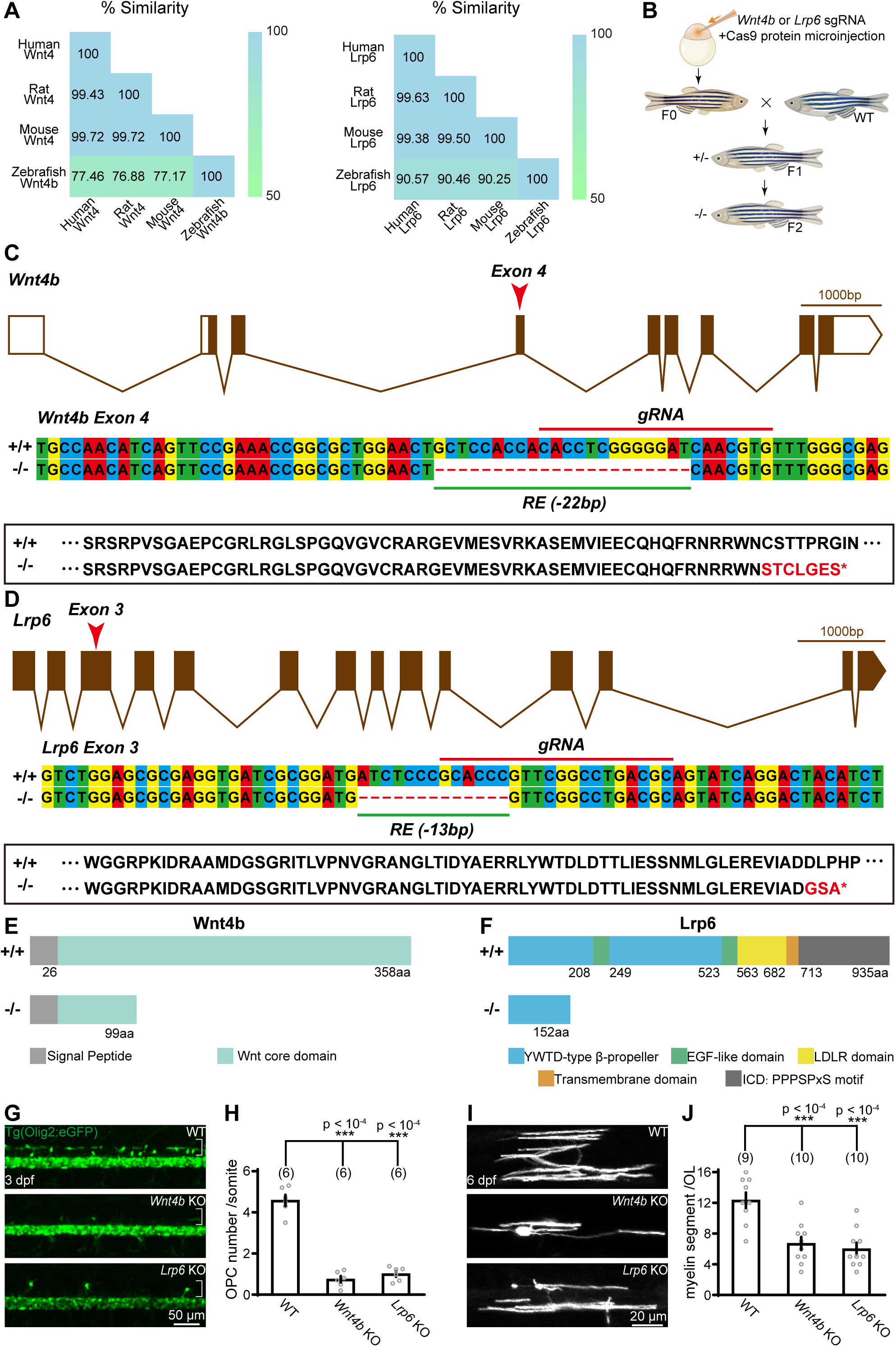
Establishment of Wnt4b and Lrp6 knockout lines. (A) Similarity of Wnt4 (left) and Lrp6 (right) proteins between zebrafish, mouse, rat and human. (B) Generation of *Wnt4b* and *Lrp6* KO fish by CRISPR/Cas9 technology. Cas9 protein and the sgRNA were microinjected into the animal pole of one-cell stage fish embryos. Injected F0 animals were raised to adulthood and outcrossed to wild-type animals to create F1 offspring. Clutches of F1 offspring were raised to adulthood and genotyped to identify heterozygous carriers of function disrupting mutant alleles. (C) Top, *Wnt4b* gene structure composed of 9 exons. Red arrowhead marks the location of the mutation in exon 4. Middle, Wild-type and mutant nucleotide sequences spanning the mutagenesis site. The gRNA target site (red line) is labeled. Bottom, Amino acid sequence indicating that the *Wnt4b* mutation results in shift in the open reading frame leading to downstream coding for a premature stop codon (*). (D) Top, *Lrp6* gene structure composed of 14 exons. Red arrowhead marks the location of the mutation in exon 3. Middle, Wild-type and mutant nucleotide sequences spanning the mutagenesis site. The gRNA target site (red line) is labeled. Bottom, Amino acid sequence indicating that the *Lrp6* mutation results in shift in the open reading frame leading to downstream coding for a premature stop codon (*). (E and F) The protein motif of Wnt4b (E) and Lrp6 (F) in WT and KO animals. (G) Visualization of dorsal migrating OPCs in WT, *Wnt4b* KO and *Lrp6* KO larvae at 3dpf. Brackets indicate the dorsal SC. (H) Statistical analysis of dorsal migrating OPCs per somite in WT, *Wnt4b* KO and *Lrp6* KO larvae at 3dpf. The number on each column represents embryos used for statistics. (I) Projected images of myelin sheath produced by an OL in WT, *Wnt4b* KO and *Lrp6* KO larvae at 6 dpf. (J) Statistical analysis of myelin segment number produced by an individual OL in WT, *Wnt4b* KO and *Lrp6* KO larvae at 6 dpf. The number on each column represents the number of OLs used for statistics. Error bars, SEM. ***p < 0.0001 (one-way ANOVA for (H and J)).

**Figure S6.**
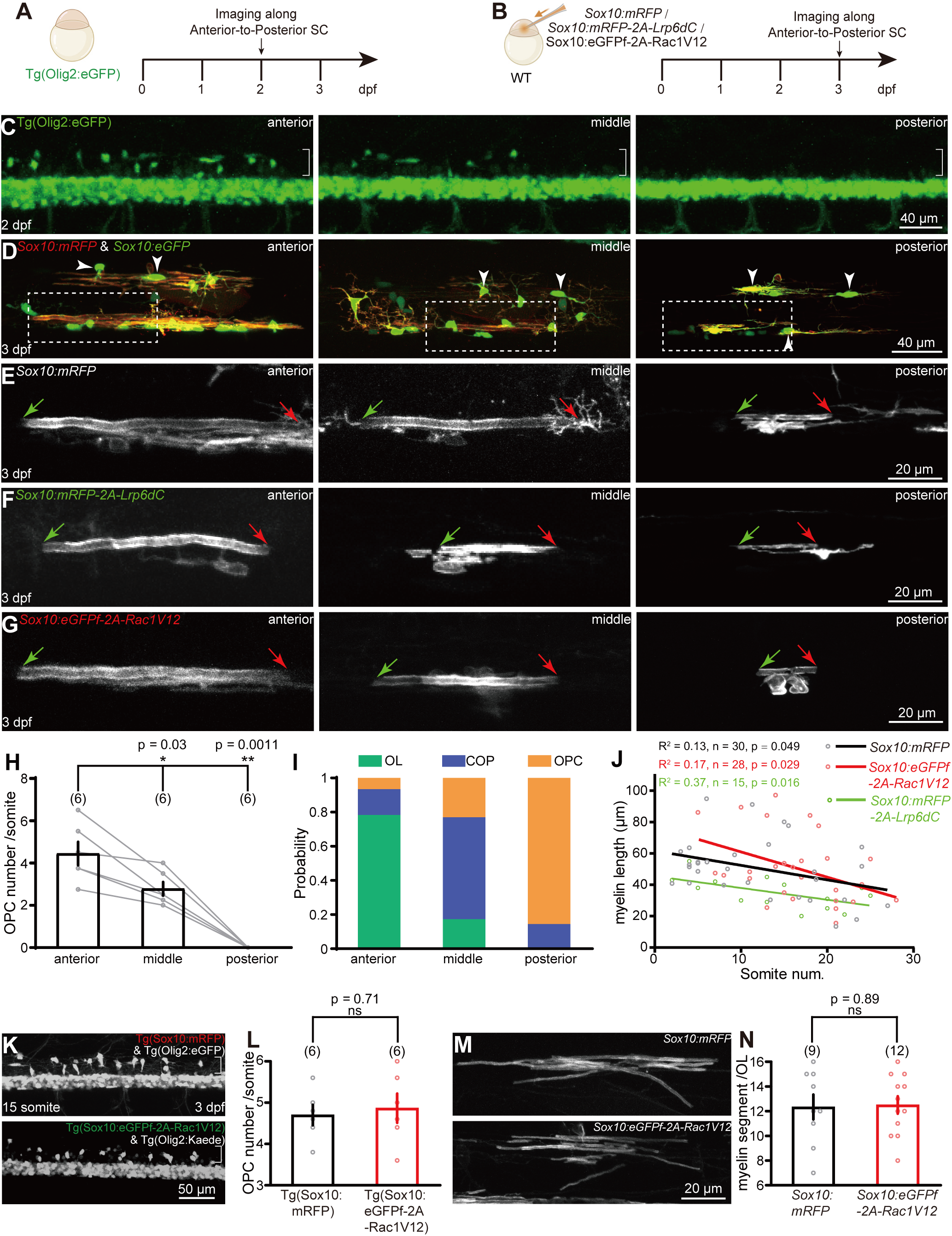
Anterior-to-posterior developmental gradient of OPC migration and OL myelination. (A) Schematic diagram displaying the imaging procedures of oligodendrocytes dorsal migration. (B) Diagram of single M-axon-wrapping myelin sheath mosaically labeled by *Sox10:mRFP*, *Sox10:mRFP-2A-Lrp6dC* and *Sox10:eGFPf-2A-Rac1V12* transient expression. Timeline indicating the imaging procedures. (C) Visualization of dorsal migrating OPCs distributed along anterior-to-posterior axis of the SC in Tg(Olig2:eGFP) larvae at 2 dpf. The white brackets indicate the dorsal SC. (D) Confocal imaging of oligodendrocyte linage cells with cytoplasm in green and membrane in red in the anterior (left), middle (middle), and posterior (right) region of the SC in 3-dpf larvae transiently expressing both *Sox10:eGFP* and *Sox10:mRFP*. Arrowheads in the left, middle, and right panels indicate myelin-formed differentiated OLs, pre-differentiated multi-process committed oligodendrocyte progenitor cells (COPs), undifferentiated bipolar OPCs, respectively. Dotted boxes in the left, middle, and right panels indicate M-axon-wrapping myelin sheath enlarged in the left, middle, and right panels in (E). (E) Visualization of M-axon-wrapping myelin distributed along anterior-to-posterior axis of the SC in *Sox10:eGFP* larvae at 3 dpf. (F) Confocal images of M-axon-wrapping myelin distributed along anterior-to-posterior axis of the SC in *Sox10:mRFP-2A-Lrp6dC* larvae at 3 dpf. (G) Confocal images of M-axon-wrapping myelin distributed along anterior-to-posterior axis of the SC in *Sox10:eGFPf-2A-Rac1V12* larvae at 3 dpf. (H) Statistical analysis of the number of dorsal migrating OPCs per somite in the anterior, middle, and posterior region of the SC at 2 dpf. Data were obtained from 6 embryos. The number on each column represents the number of embryos used for statistics. (I) Comparison of the percentage of OLs, COPs, and OPCs in the anterior, middle, and posterior region of the SC at 3 dpf. (J) Linear regression of the length of M-axon-wrapping myelin distributed along anterior-to-posterior axis of the SC in *Sox10:mRFP* (black), *Sox10:mRFP-2A-Lrp6dC* (green) and *Sox10:eGFPf-2A-Rac1V12* (red) larvae at 3 dpf. “n” represents the number of M-axon wrapping myelin used for statistics. (K) Visualization of dorsal migrating OPCs in Tg(Sox10:mRFP) & Tg(Olig2:eGFP) (top) and Tg(Sox10:eGFPf-2A-Rac1V12) & Tg(Olig2:Kaede) (bottom) larvae at 3 dpf. Tg(Olig2:Kaede) was photoconverted before imaging. Brackets indicate the dorsal SC. (L) Statistical analysis of dorsal migrating OPCs per somite in Tg(Sox10:mRFP) and Tg(Sox10:eGFPf-2A-Rac1V12) larvae at 3 dpf. The number on each column represents embryos used for statistics. (M) Projected images of myelin sheath produced by an OL in *Sox10:mRFP* (top) and *Sox10:eGFPf-2A-Rac1V12* (bottom) larvae. (N) Statistical analysis of myelin segment number produced by an individual OL in *Sox10:mRFP* and *Sox10:eGFPf-2A-Rac1V12* larvae. The number on each column represents the number of OLs used for statistics. Error bars, SEM. ns, not significant; *p < 0.05, **p < 0.01 (two-tailed unpaired Student’s t-test for (L) and (N); one-way ANOVA for (H)).

**Figure S7.**
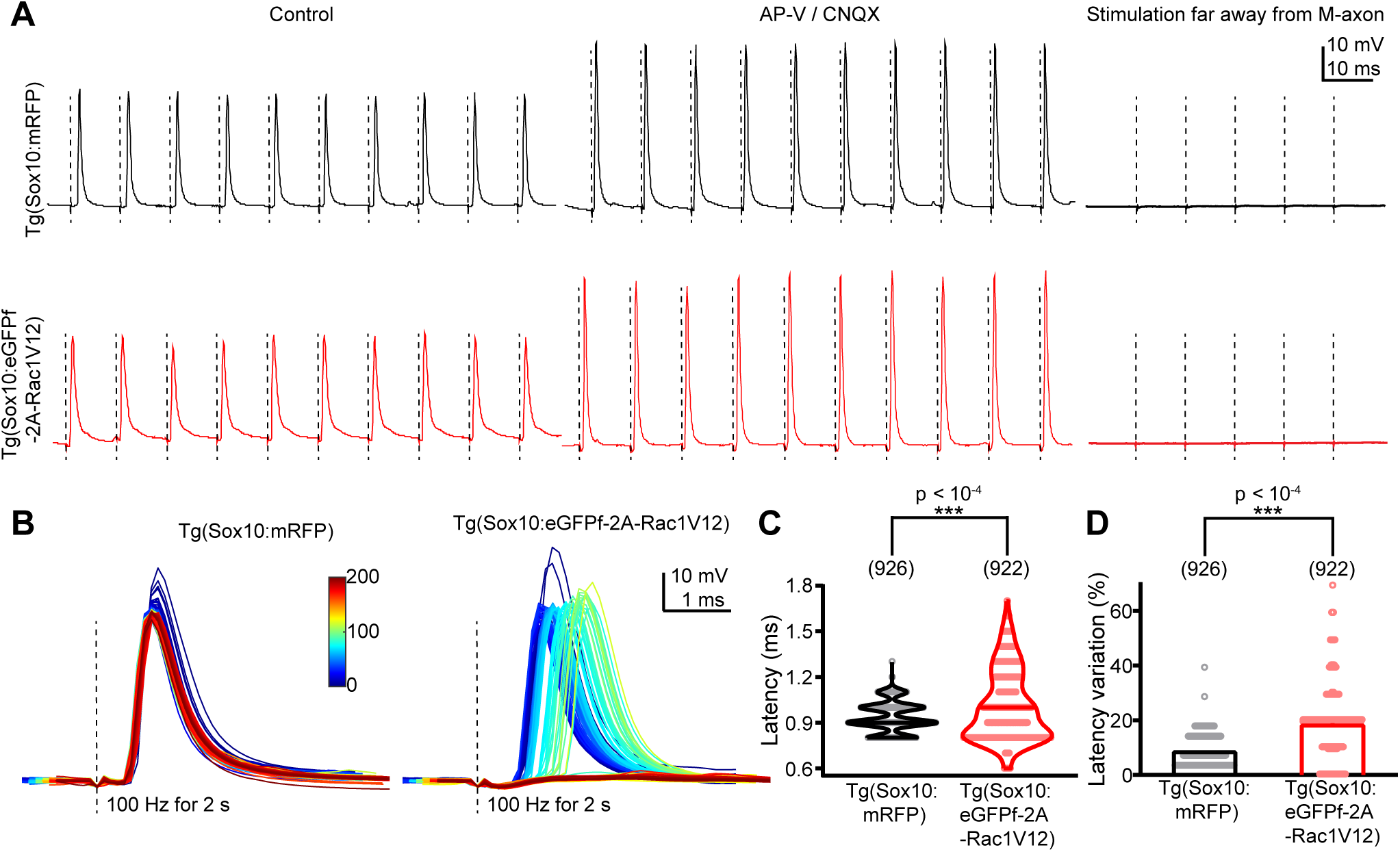
Disruption of anterior extension preference impairs the faithful transduction of APs along M-axons. (A) Samples of APs evoked by electrical stimulation (100 Hz, 100 ms) of M-axons in Tg(Sox10:mRFP) (top) and Tg(Sox10:eGFPf-2A-Rac1V12) (bottom) larvae under control (left), co-application of AP-V (100 μM) and CNQX (50 μM) (middle), and when the stimulating electrode was far away from M-axon (right). Dotted lines indicate the timing of electrical stimulation. (B) Representatives of APs evoked by electrical stimulation (100 Hz, 2 s) of M-axons in Tg(Sox10:mRFP) (left) and Tg(Sox10:eGFPf-2A-Rac1V12) (right) larvae. Dotted lines indicate the timing of electrical stimulation. Blue to red colors indicate the first to last evoked APs. (C) Latency of APs evoked by electrical stimulation (100 Hz, 2 s) of M-axons in Tg(Sox10:mRFP) and Tg(Sox10:eGFPf-2A-Rac1V12) larvae. Data were obtained from 926 evoked-APs in 5 Tg(Sox10:mRFP) larvae and 922-evoked APs in 6 Tg(Sox10:eGFPf-2A-Rac1V12) larvae. The number on each column represents the number of evoked APs. (D) Latency variation of APs evoked by electrical stimulation (100 Hz, 2 s) of M-axons in Tg(Sox10:mRFP) and Tg(Sox10:eGFPf-2A-Rac1V12) larvae. Same data set with (C) was used. The number on each column represents the number of APs used for statistics. Error bars, SEM. ***p < 0.001 (two-tailed unpaired Student’s t-test for (C) and (D)).

